# Spatial transcriptomics from pancreas and local draining lymph node tissue reveals a lymphotoxin-β signature in human type 1 diabetes

**DOI:** 10.1101/2025.05.19.654940

**Authors:** Miguel A. Medina-Serpas, Maigan Brusko, Gregory J. Golden, Martha Campbell-Thompson, Trevor Rogers, Amanda L. Posgai, Eline T. Luning Prak, Chengyang Liu, Klaus H. Kaestner, Ali Naji, Michael R. Betts, Lauren M. McIntyre, Mark A. Atkinson, Todd M. Brusko

## Abstract

This study explores the inflammatory response observed in pancreata and pancreatic lymph node (pLN) samples obtained throughout the natural history of type 1 diabetes (T1D) including non-diabetic individuals and non-diabetic autoantibody positive individuals with high susceptibility using spatial transcriptomics (ST). Integration of ST with public single-cell RNA sequencing data enabled interrogation of transcriptional alterations in T1D pathogenesis across both tissues and cellular scales. In the T1D pancreas, we observed global upregulation of multiple inflammation-associated transcripts, including regenerating islet-derived (*REG*) family genes, complement factor 3 (*C3*), *SOD2*, and *OLFM4*, and highlighted cellular candidates potentially contributing to these signatures. Within the T1D pLN, we observed spatially restricted upregulation of lymphotoxin-β (*LTB*) alongside follicular dendritic cell (FDC)-associated transcripts including *FDCSP*, *CLU*, and *FCER2*. Collectively, these findings highlight distinct inflammation signatures in the pancreas and regional pLN which can help inform the development of future therapeutic interventions.

## INTRODUCTION

Recent analyses of the human pancreas and local pancreatic lymph nodes (pLN) acquired from type 1 diabetic (T1D) organ donors, have addressed a series of major knowledge voids regarding the cellular frequencies, phenotypes, and autoreactive lymphocyte receptor repertoire signatures associated with T1D pathogenesis^1–5^. More specifically, consortia-based efforts, including the Network of Pancreatic Organ donors with Diabetes (nPOD)^6^ and the Human Pancreas Analysis Programs (HPAP)^7^, have allowed for vital and unique access to these target tissues from organ donors with T1D, non-diabetic individuals, and those at elevated risk for developing disease (i.e., autoantibody positive, non-diabetic persons). Such efforts have dramatically improved our ability to address a series of questions regarding the tissue-specific autoimmunity that occurs during the natural history of T1D^8^. However, there remains a critical need to identify therapeutically targetable pathways in individuals with failed peripheral tolerance to enable future disease prevention efforts.

High-throughput single-cell profiling represents a powerful method for assessing cellular heterogeneity within tissues by identifying cell-type specific molecular profiles suspected to contribute to the histopathological events observed in T1D or other disorders^3,9–11^. Despite their utility, one notable limitation for these techniques involves the potential introduction of confounders inherent to the generation of single-cell suspensions, including destruction of the tissue architecture, loss of stromal and rare cell types, and possible alterations in cellular phenotypes^12–14^. Sequencing-based spatial transcriptomic (ST) techniques affords the ability to address this challenge by allowing for gene expression quantitation in whole tissue sections at variable scales (i.e., multicell vs. single-cell vs. subcellular) and gene detection sensitivity (unbiased whole transcriptome vs. a pre-defined gene panel)^15^. Furthermore, ST methods utilizing transcript-specific probe sets for gene detection exhibit increased sensitivity by overcoming stochastic dropout observed in single-cell RNA-sequencing (scRNA-seq) based approaches^16^. These techniques are also compatible with analysis of fixed tissues that inherently preserve the underlying tissue microenvironment, enabling simultaneous consideration of individual cellular positions, cellular neighborhoods, cell-cell contacts, and gene expression^17^.

Herein, we performed multi-cell resolution ST profiling on formalin-fixed paraffin-embedded (FFPE) pancreata and donor-matched pLN sections matched to the same pancreatic anatomical region (i.e., head, body, or tail) acquired from human organ donors in the HPAP collection^7^. This cross-sectional cohort represents donors across the natural clinical history of T1D, including non-diabetic autoantibody negative (AAb-) controls (ND), non-diabetic donors with 1 islet AAb (sAAb+), non-diabetic donors with ≥2 islet AAb (mAAb+), and T1D donors. Furthermore, we leveraged annotated pancreas^18^ and pLN^3^ scRNA-seq datasets for integration and cell deconvolution, enabling the assessment of relative changes in cellular frequencies across both tissues. From this analysis, we identified significant immune cell enrichment in the pancreas of at-risk donors and alterations in immune cell subset enrichment in the pLN across T1D disease states. The data reported herein provide evidence for dysregulated gene profiles in distinct pancreatic regions, as well as innate and adaptive inflammatory signatures in the pLN, along with candidate biomarkers to guide future interventional therapies for T1D.

## RESULTS

### Interrogation of matched pancreas and pLN by spatial transcriptomics

To gain insights into the tissue-specific inflammatory events resident to the human pancreas and pLN across the natural history of T1D, we performed ST profiling at 55 µM capture spot resolution on FFPE pancreatic tissue sections from 20 human organ donors, including: ND control donors (n=5), sAAb+ donors without diabetes (n=3), mAAb+ donors without diabetes (n=4), and donors with a clinical history of T1D (n=8; mean disease duration=4.19 years ± 2.85). For these studies, pLN sections were acquired from the same pancreatic donors when available (16/20). Per-donor tissue availability, islet-AAb status, and T1D-associated HLA genetic risk data are summarized in **Figure 1A** with detailed donor characteristics provided in **Supplemental Table 1**.

**Figure 1:**
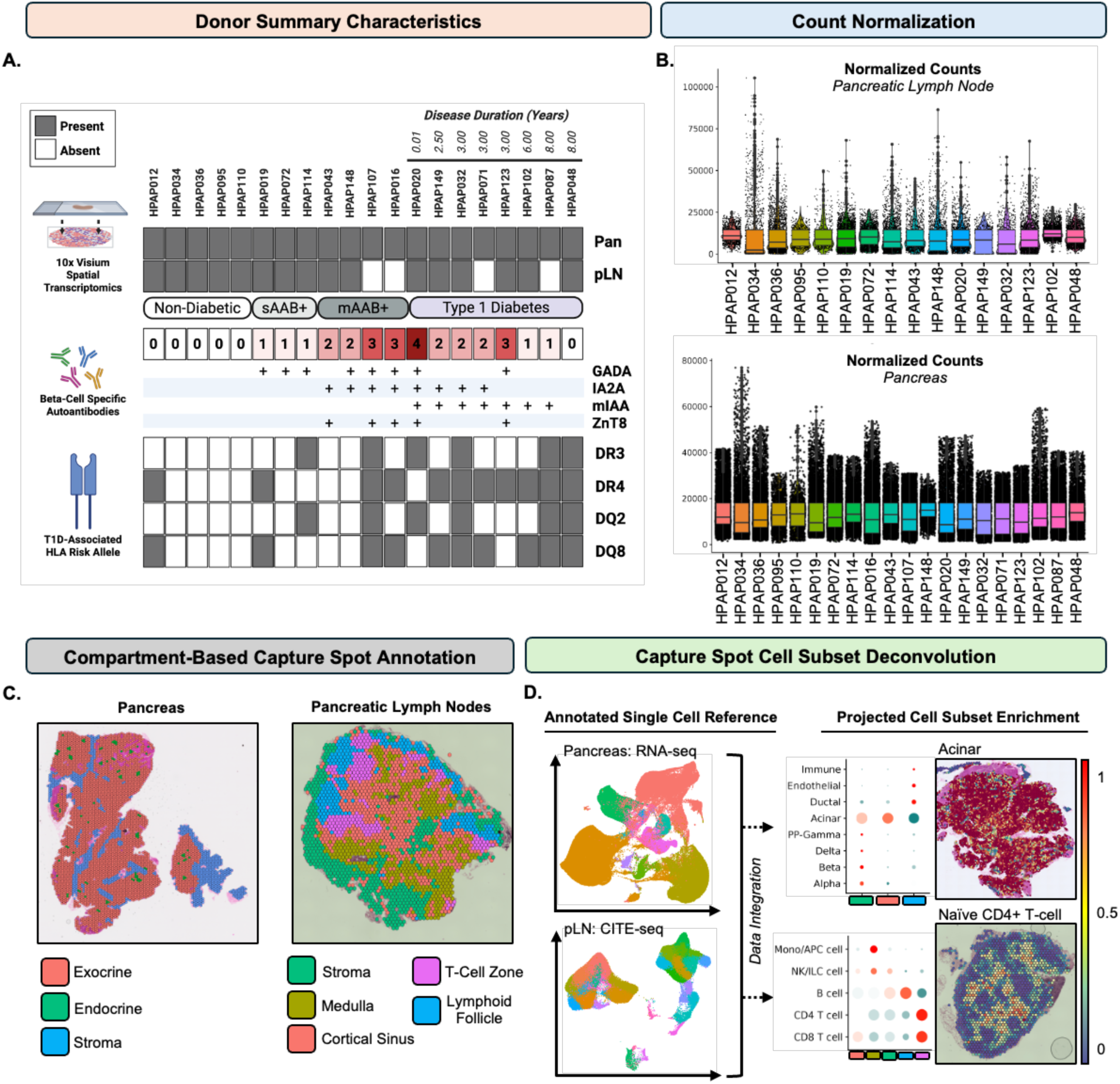
Summary of donor characteristics and spatial transcriptomics workflow. (A) Summary of individual donor characteristics including assayed tissue types, number of islet specific autoantibodies, and T1D risk associated HLA diplotype. (B) Violin boxplots depicting normalized read counts across individual donors for pLN (top) and pancreas (bottom). (C) Representative pancreas (left) and pLN (right) sections depicting spatial arrangement of annotated tissue regions. (D) Cell deconvolution of pancreas and pLN spatial data sets was performed via anchor-based CCA using annotated single cell gene expression reference derived from the same tissue type. This method yields proportion values (0.0 – 1.0) at each capture spot position corresponding to the expected proportional enrichment of all cell types represented in the annotated single cell reference.

For each tissue, capture spots were integrated into a latent space, via anchor-based canonical correlation analysis (CCA),^19^ yielding two separate pancreas and pLN datasets containing 113,356 capture spots for pancreas and 25,584 total capture spots for pLN. To address inter-donor variability of unnormalized count distributions following integration (**Supp. Fig. 1A & 1B**), we applied an upper quartile normalization strategy^20^ across each dataset (**Fig. 1B**). We quantitatively (**Supp. Fig. 1C & 1D**) and qualitatively (**Supp. Fig. 1E & 1F**) assessed the integration quality and observed comparable mean embedding values across donors and representation across disease status.

We next annotated capture spots by their tissue compartment localization for each respective tissue using histopathological features (**Supp. Table 2 & 3**) identified in the hematoxylin and eosin (H&E) stained sections prepared during spatial expression profiling. Expression of canonical marker genes corresponding to known cell types also aided in capture spot annotation (**Fig. 1C**). At 55µm diameter, capture spots reflect expression profiles derived from multiple cells and/or cell types. We thus applied *in silico* cell deconvolution^21^ to calculate the predicted proportion of constituent cell types within capture spots using single-cell gene expression reference data^3,22^ (isolated pancreatic islets – scRNA-seq [https://hpap.pmacs.upenn.edu^18^]; pLN – CITE-seq [GSE221787]) (**Fig 1D, Supp. Fig. 1G & 1H**). Donor overlaps between our ST cohort and the respective pancreas and pLN single-cell reference is outlined in **Supp. Table 1**.

### Differentially expressed genes (DEGs) across defined compartments of the pancreas

Marker gene analysis of annotated pancreas compartments identified enrichment of canonical endocrine (*GCG -* α-cell; *INS* -β-cell; *SST* - δ-cell), exocrine (*CELA2A, CELA3B*, *PRRS3)*, and stromal (*FN1*, *ACTA2*, *MYH11*) genes within the expected annotated compartments (**Fig. 2A**). Corresponding whole-slide scanned H&E-stained pancreas sections were examined to ensure the tissue gene expression signature matched the underlying histology (**Fig. 2B**). Principal component analysis (PCA) was used to visualize the expression profiles for each compartment (**Fig. 2C**). We then performed within compartment differential expression testing between AAb+ (sAAb+ and mAAb+ considered together for increased statistical power), T1D, and ND individuals (**Table S4-6**). 28 DEGs showed evidence of compartment specific expression across all comparisons with the majority restricted to the endocrine compartment (17/28, 61%) (**Fig. 2D**). Similarly, the endocrine compartment accounted for the greatest number of DEGs overall (n = 60), followed by the exocrine (n = 51) and stromal compartments (n = 28) (**Fig. 2E**).

**Figure 2:**
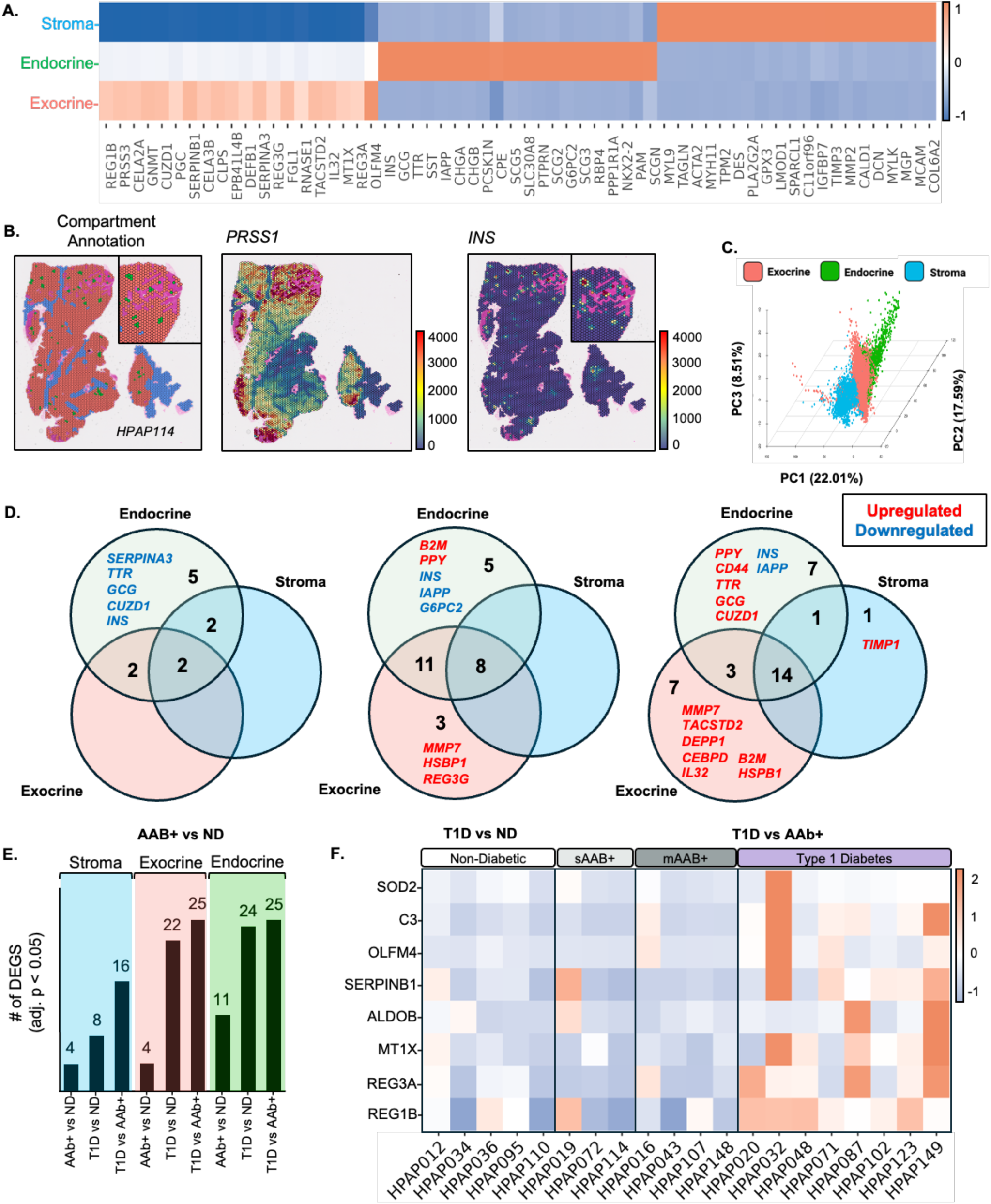
Compartment-level differential expression testing in pancreas reveals distinct spatially restricted and tissue-wide inflammation signatures. (A) Heatmap depicting the relative scaled expression of the top 20 marker genes corresponding to the exocrine, endocrine, and stroma compartments in donor pancreas. (B) Representative spatial mapping of annotated pancreas compartments with marker gene expression for both the exocrine and endocrine compartments plotted spatially. (C) Projection of all pancreas capture spots (113,356) onto 3-dimensional principal component plot. (D) Bar graph depicting the number of differentially expressed genes detected in each annotated compartment across 3 pairwise comparisons (AAB+ vs ND; T1D vs ND; T1D vs AAB+). (E) Venn diagram plots indicating the number of DEGs detected between annotated pancreas compartments across 3 pairwise comparisons. Refer to Supp. Table 4-6 for individual genes, Log_2_FC difference, and significance values. (F) Heatmap depicting the relative scaled expression of significant DEGs detected across multiple tissued compartments in T1D sections relative to ND and non-diabetic AAb+ donors across individual donors.

### Endocrine compartment-specific DEGs indicate reduced **β**-cell function, increased inflammation, and tissue injury in AAb+ and T1D pancreas

Within the endocrine compartment, we observed down regulation of *INS, GCG, TTR, CUZD1,* and *SERPINA2* among AAb+ donors relative to ND (**Fig. 2E & Supp. Fig. 2B**). Reduced *IN*S expression may indicate declining β-cell mass or function prior to T1D onset which is consistent with prior reports^23,24^. Similarly, reduced *GCG* expression may also point towards α-cell dysfunction which has been reported in T1D^25^ and in individuals with a single GAD AAb where it was linked to altered expression of oxidative phosphorylation genes^26^. *TTR* (transthyretin) encodes a transport protein highly expressed in α-cells but also functions in insulin release and protection of β-cells from apoptosis^27^. CUZD1 (CUB and zona pellucida-like domain-containing protein 1) is reported as reduced in pancreatitis and is an autoantibody target in Crohn’s disease^28^. SERPINA3 (alpha-1-antichymotrypsin) is an acute-phase protein with purported roles in diabetic nephropathy and heart disease^29,30^. Collectively, these gene expression differences point towards aberrations in β-cell function and potentially, inflammation within endocrine areas in AAb+ individuals.

The endocrine compartment of T1D subjects exhibited reduced expression of β-cell-enriched transcripts *INS*, *IAPP* (islet amyloid polypeptide), and *G6PC2* (glucose-6-phosphatase catalytic subunit 2), as expected relative to ND controls. *PPY* (pancreatic polypeptide) expression was also increased in T1D, possibly due to a minor imbalance of pancreatic head region sections (**Supp. Table 1**), as cells producing this hormone are found primarily in the ventral lobe of the pancreatic head^31,32^. Although direct detection of HLA receptor transcripts was not possible with the Visium CytAssist assay due to their allelic hypervariability, we observed upregulation of the class I HLA subunit *B2M (*β2 microglobulin) in T1D subjects relative to ND (**Fig. 2E & Supp. Fig. 2B),** as reported by others^33–35^. We also observed upregulation of the cell adhesion and migration gene *CD44* in the T1D endocrine compartment relative to AAb+ donors (**Fig. 2E & Supp. Fig. 2C**), potentially indicative of tissue injury^36^.

### Exocrine compartment changes support chronic pancreatic inflammation in T1D

We detected upregulation of *MMP7* (matrix metalloproteinase 7)*, HSBP1* (heat shock factor binding protein 1), and *REG3G* (regenerating islet-derived protein 3 gamma) in T1D sections relative to ND controls (**Fig. 2E & Supp. Fig. 2D, E**). These genes have been implicated in models of acute pancreatitis: *MMP7*^37^ and *REG3G*^38^ are thought to contribute to inflammation, whereas *HSBP1*^39^ has been shown to have protective qualities. Furthermore, we also detected upregulation of *TACSTD2, DEPP1, CEBPD, IL32,* and *B2M* in the T1D exocrine compartment when compared to AAb+ donors (**Fig. 2E & Supp. Fig. 2F**). Exocrine-specific upregulation of *B2M* is supported by immunohistochemical staining which revealed expression of class I HLA protein on the surface of acinar cells in recent-onset T1D cases but not in non-diabetic controls^40^. Our observation in AAb+ individuals suggest inflammation of the exocrine pancreas may exist before clinical onset. Lastly, we observed altered expression of *TACSTD2*^41^*, DEPP1*^42^*, CEBPD*^43^, and *IL32*^44^ in the exocrine pancreas which has been associated with pancreatic malignancies, however, in this context they may suggest a chronic inflammatory environment in the T1D pancreas.

### DEGs shared across compartments indicate a global inflammatory signature in T1D pancreas

Among T1D sections, we observed global upregulation of additional REG gene family members (*REG1B, REG3A*), as well as *MT1X* (Metallothionein 1X), *ALDOB* (Aldolase B), *SERPINB1* (leukocyte elastase inhibitor), *OLFM4* (olfactomedin 4)*, C3* (complement component 3), and *SOD2* (superoxide dismutase 2) (**Fig. 2F**). Relative to ND, REG genes were significantly downregulated in AAb+ donors, yet significantly upregulated in T1D, with relatively consistent expression patterns visualized across individual donors in these groups (**Fig. 2F**). This expression pattern may suggest that REG genes are variable across stages and dynamically regulated by inflammation during T1D.

*MT1X*, which was upregulated in T1D versus AAb+ and/or ND across all three tissue compartments (**Fig. S2**), is a negative regulator of insulin secretion that is primarily expressed by β-cells but associates with β-cell failure with increased expression reported in type 2 diabetes (T2D) islets^45^. We also observed increased expression of *ALDOB* in T1D endocrine and exocrine tissue regions as compared to both the ND and AAb+ groups. *ALDOB* is also primarily expressed by β-cells and its induction negatively correlates with insulin secretion^46^. Together, these observations likely reflect stress in residual β-cells after T1D onset.

Functionally, *SERPINB1, OLFM4, C3,* and *SOD2* are involved in both innate and adaptive immunity^47–49^, therefore, we assessed the expression of these immune-related genes in the pLN CITE-seq reference dataset (GSE221787) to infer the immune cell type composition that may contribute to this signature in the pancreas. *C3* and *SOD2* appeared to be generally enriched across myeloid cell subsets (**Supp. Fig. 2J**). *OLFM4* and *SERPINB1* were expressed across multiple putatively annotated cell types, including various distinct memory B-cell subsets. However, only *OLFM4* was found to be expressed across naïve and central memory CD4^+^ T-cell subsets and NK/innate lymphoid cells (ILCs), whereas only *SERPINB1* was found to be enriched in myeloid populations (**Supp. Fig. 2J**). These data indicate the heightened pancreatic immune activity in T1D, even in the absence of observable insulitis by H&E pathology analysis (**Supp. Table 3**).

### Immune enrichment in mAAb+ pancreas prior to T1D onset

Next, in order to characterize inflamed regions of the pancreas at the cellular level, we used an independently prepared scRNA-seq reference prepared from isolated human pancreatic islets^18^ to deconvolute immune-enriched capture spots. Prior to cell deconvolution, we removed all cells acquired from T2D subjects in the scRNA-seq reference. The reference includes annotations for major endocrine (α, β, δ-cells) and exocrine (acinar, ductal) cell types, with immune cells represented as a singular heterogeneous population. Spatial mapping revealed localization of acinar, endocrine, ductal, and epithelial cell types consistent with the compartmental annotation (**Supp. Fig. 3A-B**). T1D donors exhibited minimal immune cell enrichment in the endocrine compartment, consistent with histopathological examination of paired H&E images, which only identified insulitis in a single donor (**Supp. Table 3**). Instead, immune cell signatures were enriched on average in the stromal compartment of the pancreas among all donor groups (**Supp. Fig. 3A**). Accordingly, instances of localized mononuclear cell infiltration were found along interlobular and fascial connective tissues in cases described as chronically inflamed after histopathological assessment (**Supp. Table 3**). The presence of mononuclear cells in immune-enriched regions was visually confirmed (**Fig. 3A**). We then evaluated the predicted immune cell enrichment as calculated by cell deconvolution for differences across disease status after removing all capture spots predicted to contain no immune cell presence. We observed mAAb+ donors had significantly higher immune proportion values relative to all other donor groups (**Fig. 3B**).

**Figure 3:**
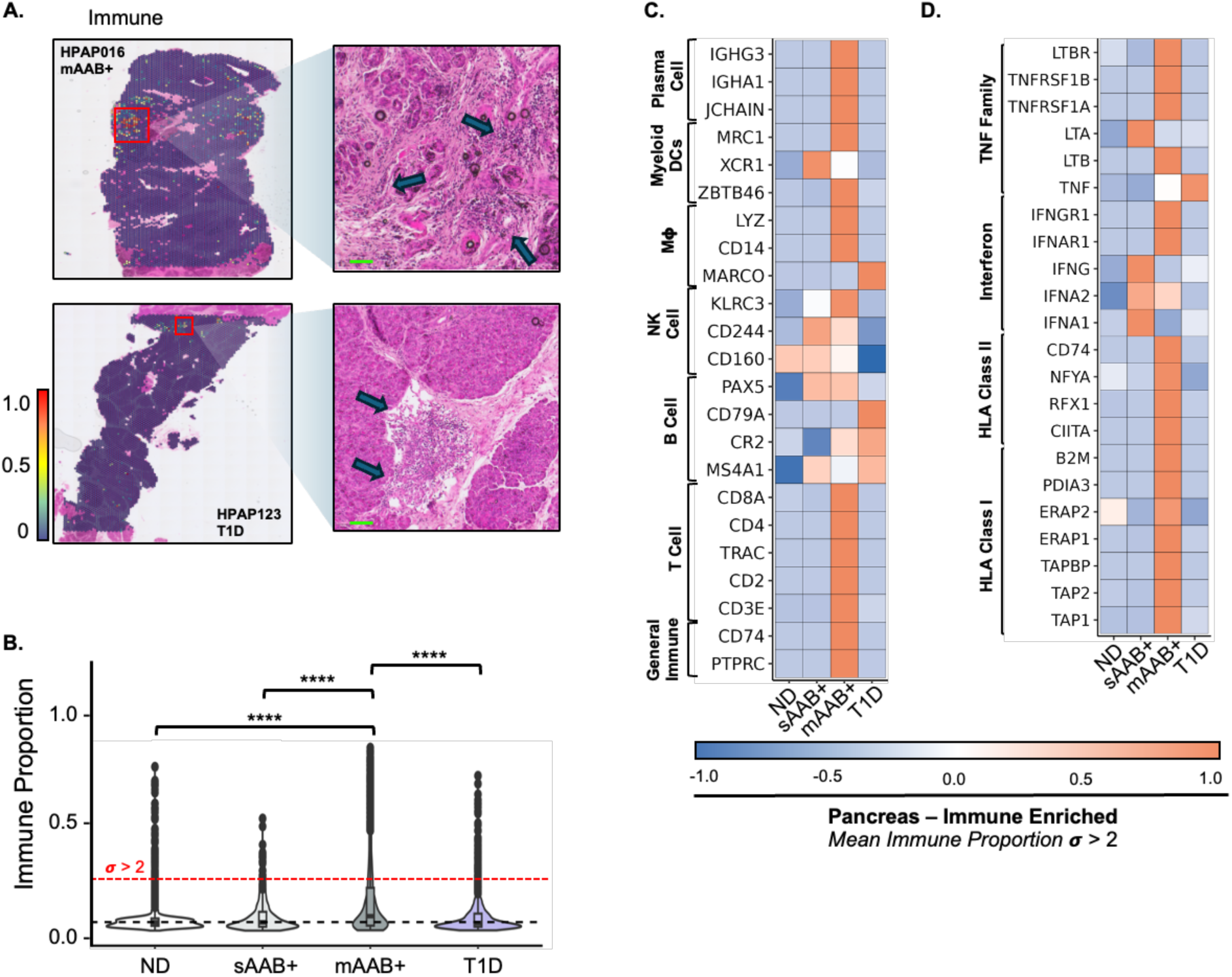
Global assessment of the pancreas unveils a distinct immune and inflammation associated signature preceding the development of clinical T1D. (A) Unbiased identification of immune cell containing capture spots by cell deconvolution in representative pancreas sections (left). Regions predicted to contain high immune cell enrichment were followed up by examination of corresponding H&E stained sections which revealed the presence of mononuclear cell infiltration (right; denoted by arrows). (B) Violin-boxplot comparing the level of immune cell enrichment in capture spots calculated by cell deconvolution across disease status. ND: non-diabetic AAb-; sAAb+: non-diabetic single AAb+; mAAB+: non-diabetic multiple AAb+; T1D: type 1 diabetes. Unpaired students T-test. Significance denoted by *p. value < 0.05, **p. value < 0.005, *** p. value < 0.0005, and ****p. value < 0.00005. The red line along the y-axis represents the level of immune enrichment 2 standard deviations (α, 0.21) above the non-diabetic mean. Boxplot edges denote the lower and upper quartile with the center dash representing the mean. Dots represent outliers. (C) Heatmap depicting the relative scaled expression of general immune and canonical T cell, B cell, NK cell, macrophage (Mχτ), myeloid dendritic cells, and plasma cell marker genes in capture spots identified as immune enriched. (D) Heatmap depicting the relative scaled expression of established class I and class II antigen presentation, interferon, and TNF signaling genes within capture spots identified as immune enriched.

Next, we measured the expression of canonical immune cell type marker genes within immune cell containing capture spots to determine their cellular composition and compared across status. We directed our focus on positions considered immune enriched. This was defined as capture spots possessing an immune proportion ≥ 2 standard deviations (σ) above the ND mean which is representative of heightened immune activity, potentially in response of localized inflammation. We then visualized the expression levels of known marker genes expressed across a variety of immune cell types (*PTPRC, CD74*) and those specific or enriched in T cells (*CD3E, CD2, TRAC, CD4, CD8A*), B cells (*MS4A1, CR2, CD79A, PAX5*), NK cells (*NCAM1, CD160, CD244, KLRC3*), monocyte/macrophages (*MARCO, CD14, CD68*), classical dendritic cells (DCs; *ZBTB46, XCR1, MRC1*), and plasma cells (*JCHAIN, IGHA1, IGHG3*) across donor groups within immune enriched capture spots (**Fig. 3C**).

Consistent with the overall greater predicted immune enrichment in mAAb+ donors (**Fig. 3B**), the highest relative expression levels of all measured immune marker genes were detected in this group (**Fig. 3C**). Except for the enrichment of some NK cell marker genes, the immune composition of sAAb+ donors were comparable to ND. T1D donors were enriched for B cell-associated transcripts rather than T cell genes.

Finally, to assess the expression of inflammatory genes previously implicated in the pathogenesis of T1D^33,50–52^, expression of interferon (*IFNG, IFNA1, IFNA2, IFNAR1, IFNGR1*), tumor necrosis family (*TNF*, *LTA, LTB, TNFRSF1A, TNFRSF1B, LTBR*), and antigen presentation genes (*TAP1, TAP2, TABP, ERAP1, ERAP2, PDIA3, B2M, CIITA, CD74*) were compared across disease states. We also included additional genes directly involved in regulating class II HLA expression *(NFYA, RFX1)*^53,54^ as surrogates. Mirroring our assessment of immune markers, expression of nearly all inflammation-associated genes were most elevated in mAAb+ donors (**Fig. 3D**). Interestingly, we also detected upregulation of *LTA, IFNG, IFNA1,* and *IFNA2* in sAAb+ donors (**Fig. 3D**). Collectively, these results indicate disease-associated inflammation in the target organ prior to T1D onset and highlight the heterogeneous cellular contribution to the immune signature among mAAb+ donors.

### Transcriptional characterization of major histological compartments of pLN

Next, we aimed to characterize the transcriptional landscape of the draining pLN by first defining gene expression signatures corresponding to five distinct regions based on tissue architecture identified in H&E images: the medulla (*CLEC4G, MARCO, STAB2*), stroma (*DES, MYH11, CNN1*), lymphoid follicles (*CXCL13, PAX5, CD19*), T cell zone (TCZ) (*CD3G, CD3E, TCF7*), and an indistinct region of mixed cell types (*FOXD4L5, PEX11A*) (**Fig. 4A**). The spatial gene expression pattern of regional marker genes supported these annotations (**Fig. 4B**), with the exception of the indistinct region which had lower overall gene counts (**Fig. 4C**). PCA analysis showed all regions except the mixed cell region were transcriptionally distinct from one another, resulting in its exclusion from further analysis (**Fig. 4D**). Lastly, we examined the capture spot frequency across each annotated region at the individual donor level and across each disease classification to assess for regional imbalances (**Fig. 4E**). The capture spot numbers and regional coverage showed variation caused by tissue specimen size and orientation at the time of embedding and sectioning, but all regions were well-represented in all specimens (**Supp. Table 2 & 3**).

**Figure 4:**
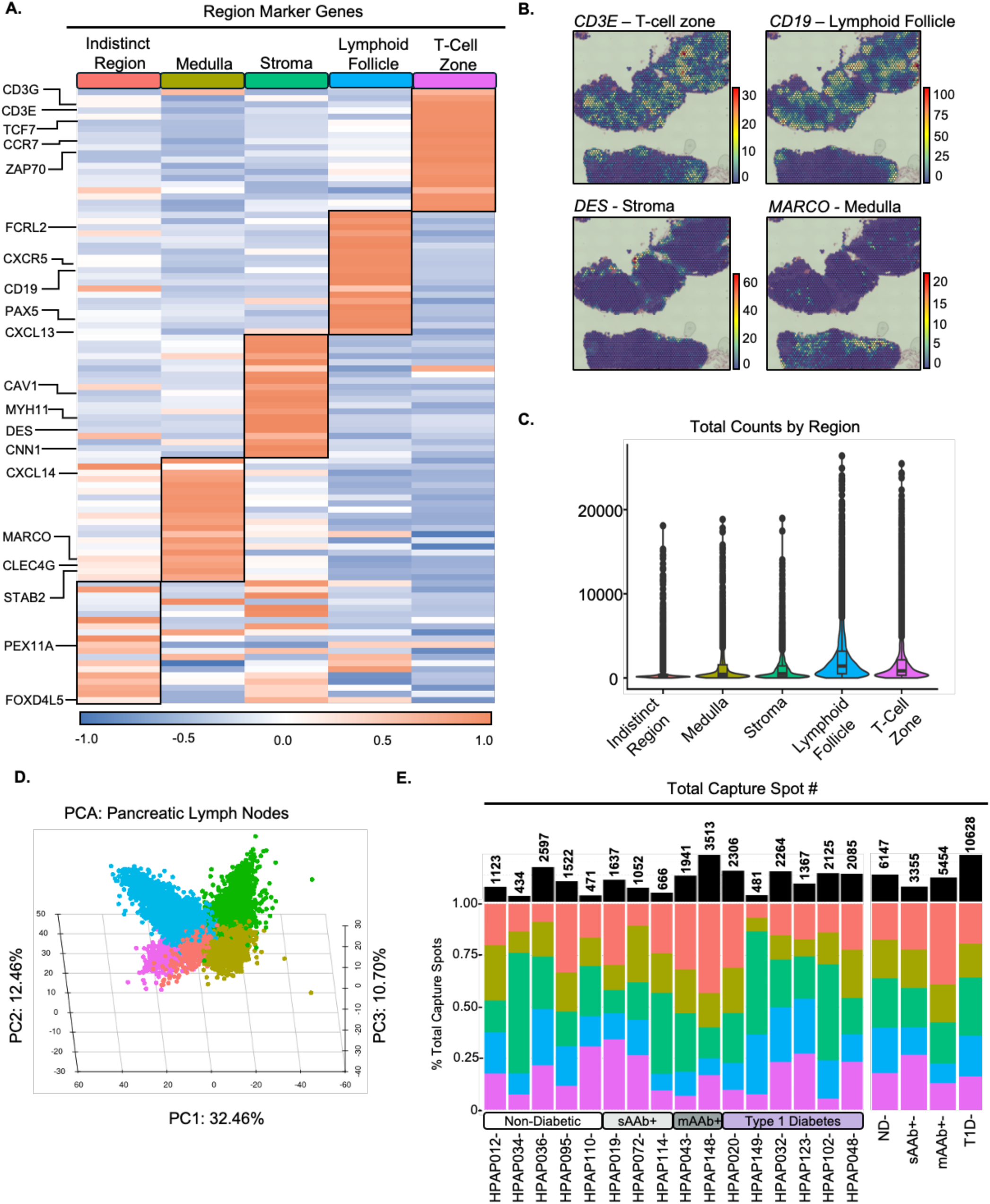
Transcriptional characterization of major histological compartments of donor pancreatic lymph nodes. (A) Heatmap depicting the relative scaled expression of the top 15 marker genes corresponding to the annotated tissue compartments in donor pLN sections. (B) Spatial gene expression profiling of representative marker genes in a representative pLN section. (C) Violin-boxplot depicting the total count distribution per annotated region in the pLN. Boxplot edges denote the lower and upper quartile with the center dash representing the mean. Dots represent outliers. (D) Projection of all pLN capture spots (25,584) onto 3-dimensional principal component plot. Colors for each point correspond to each annotated pLN region: TCZ (purple), lymphoid follicle (blue), stoma (green), medulla (gold), mixed cell region (red). (E) Stacked bar plot depicting the proportional representation of annotated pLN compartments across each donor. Colors within the stacked bar plot correspond to the annotated pLN region. TCZ (purple), lymphoid follicle (blue), stroma (green), medulla (gold), cortical sinus (red). The standard black bar plot above indicates the total number of capture spots per donor.

### Spatial mapping of immune cell heterogeneity in pLN at distinct stages of T1D

As in pancreas, we performed cell deconvolution of capture spots across all pLN regions utilizing an existing annotated CITE-seq reference (GSE221787) generated from 34 human pLN isolated at distinct stages of T1D from HPAP^3^. This reference contains 30 distinct immune subsets represented across five major cell types (CD4^+^ T cell, CD8^+^ T cell, B cell, NK/ILC, myeloid) (**Supp. Fig. 1E**). The TCZ and lymphoid follicles were predicted to be highly enriched for T and B cells, respectively, while the medulla was particularly enriched for myeloid APCs, in addition to NK cells and ILCs (**Fig. 5A**). Spatial cell type mapping of B cell and CD4^+^ T cell proportions displayed the expected spatial organization characteristic of both the lymphoid follicles and TCZ^55^ (representative sections, **Fig. 5B**).

**Figure 5:**
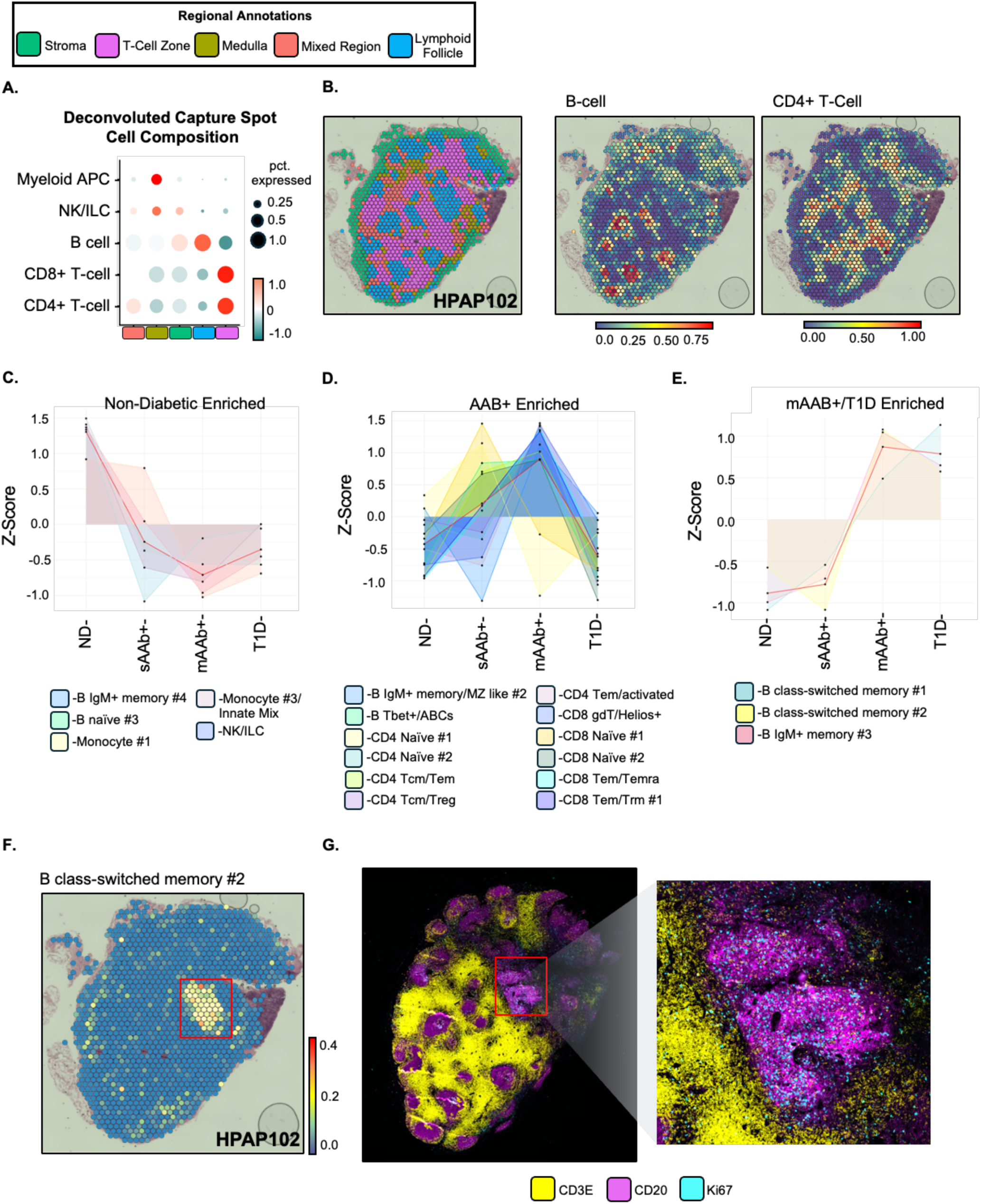
Cell deconvolution enables thorough assessment and spatial mapping of immune cell heterogeneity in donor pancreatic lymph nodes at distinct stages of T1D. (A) Dot plot depicting the relative enrichment of 5 major immune cell types across annotated pLN regions as calculated by cell deconvolution. Dot size represents the percent of total cell-type enrichment within each compartment. (B) Spatial projection of annotated tissue regions in a representative donor section alongside the relative cell type enrichment as calculated by cell deconvolution of B cell and CD4+ T cells in the same representative section. (C) Cell-type enrichment modules corresponding to immune cell identities found to be only enriched in ND sections, (D) non-diabetic AAb+ sections, and (E) in both non-diabetic mAAb+ and T1D sections. (F) Spatial enrichment profiling of B class-switched memory #2 population in representative donor pLN section. (G) 3 color immunofluorescence image of sequentially prepared donor pLN section against CD3E (yellow), CD20 (magenta), and Ki67 (teal). Imaged at 20x magnification.

Next, we examined the relative predicted immune cell composition of each pLN capture spot at the expanded 30 immune subset resolution across disease states (individual donors shown in **Supp. Fig. 4A**). To assess overall enrichment patterns across the natural history of T1D progression, cell subset enrichment scores for all capture spots were scaled, compared across disease status, and organized into four distinct modules of peak relative enrichment [modules = ND enriched, AAb+ enriched, T1D enriched, and mAAb+/T1D enriched] (**Fig. 5C-E, Supp. Fig. 4B**). Among ND donors, we observed peak enrichment of naïve B cell subsets, myeloid, and NK cells relative to all other groups (**Fig. 5C**). Interestingly, both naïve and memory/effector T cell subsets were predicted to be most enriched in AAb+ donors (**Fig. 5D**), with naive T cell subset enrichment in sAAb+ donors, and memory/effector subsets enriched in mAAb+ donors. In the pLNs of T1D donors, however, expression programs associated with distinct memory B cell subsets were heightened.

Evaluation of the mAAb+/T1D enriched module, which consists of cell types initially enriched before disease onset, contained two distinct class-switched memory (CSM) B cell subsets and an IgM+ memory subset (**Fig. 5E**). At the per-donor level, 2/2 (100%) mAAb+ and 3/6 (50%) T1D donor pLNs were identified as globally enriched for this B CSM #2 signature (**Supp. Fig. 4A**); however, after examining the spatial organization across donors, a single T1D donor (HPAP102) possessed a focal enrichment of this cell signature within an annotated lymphoid follicle region (**Fig. 5F**). We prepared a serial section of HPAP102 pLN for three-color immunofluorescence imaging against CD3E (T-cell), CD20 (B-cell), and Ki67 (proliferation) and noted extensive Ki67+CD20+ co-staining with colocalization of CD3E+ cells within this same follicle region (**Fig. 5G**). To understand the functional identity of the B CSM #2 subset, we measured the surface protein expression of memory (CD27), class-switch (IGHM), and costimulatory markers (CD40, FAS, CD86) alongside their corresponding transcripts and *BCL6*, which encodes the master regulator of germinal center (GC) B cell responses^56^, in the pLN CITE-seq reference data^3^ (**Supp. Fig. 4C-E**). We noted that B CSM #2 cells exhibited the highest surface expression of costimulatory proteins, low levels of IgM and IgD (suggestive of class-switching) and high expression of *BCL6,* suggestive of a GC-like phenotype, which is further supported by the immunofluorescence staining (**Fig. 5G**). Collectively, these findings highlight a common skewing of T cells and B cells toward effector and memory signatures in pLN of mAAb+ and T1D donors when compared to ND.

### Follicular dendritic cell (FDC) and lymphotoxin β (LTβ) responsiveness genes are enriched in AAb+ and T1D pLN

To assess alterations in gene expression across disease groups, we implemented differential expression testing of the four pLN tissue compartments – the lymphoid follicles, TCZ, stroma, and medulla. Approximately 70% (59/85) of all reported DEGs were localized to the lymphoid follicles (33/85; 38.8%) and the medulla (26/85; 30.6%) across all pairwise comparisons, whereas the fewest were associated with the TCZ (9/85; 10.5%) (**Supp. Fig. 5A**). We further noted that the majority of compartment-specific DEGs localized to the T1D lymphoid follicles relative to ND and/or AAb+ donors, suggesting altered follicular activity as a feature of disease (**Supp. Fig. 6B**). All significant DEGs across tested pLN regions were reported in **Supplemental Figure 5C-K** and **Supplemental Table 7-10**.

Within AAb+ lymphoid follicles, we observed reduced expression of the immunoregulatory gene *CD52*^57^ and the immunoregulatory purinergic receptor *P2RX5*^58^ relative to ND controls (**Fig. 6A**). We also observed reduced expression of immunoglobulin genes *IGHA1* and *JCHAIN* alongside the oxidative stress gene *TXNDC5*^59^ in AAb+ vs ND lymphoid follicles. Interestingly, differential expression testing identified the TNF-family ligand *LTB* as the most highly upregulated gene in the T1D lymphoid follicles relative to ND and AAb+ donors (**Fig. 6B-C**).

**Figure 6:**
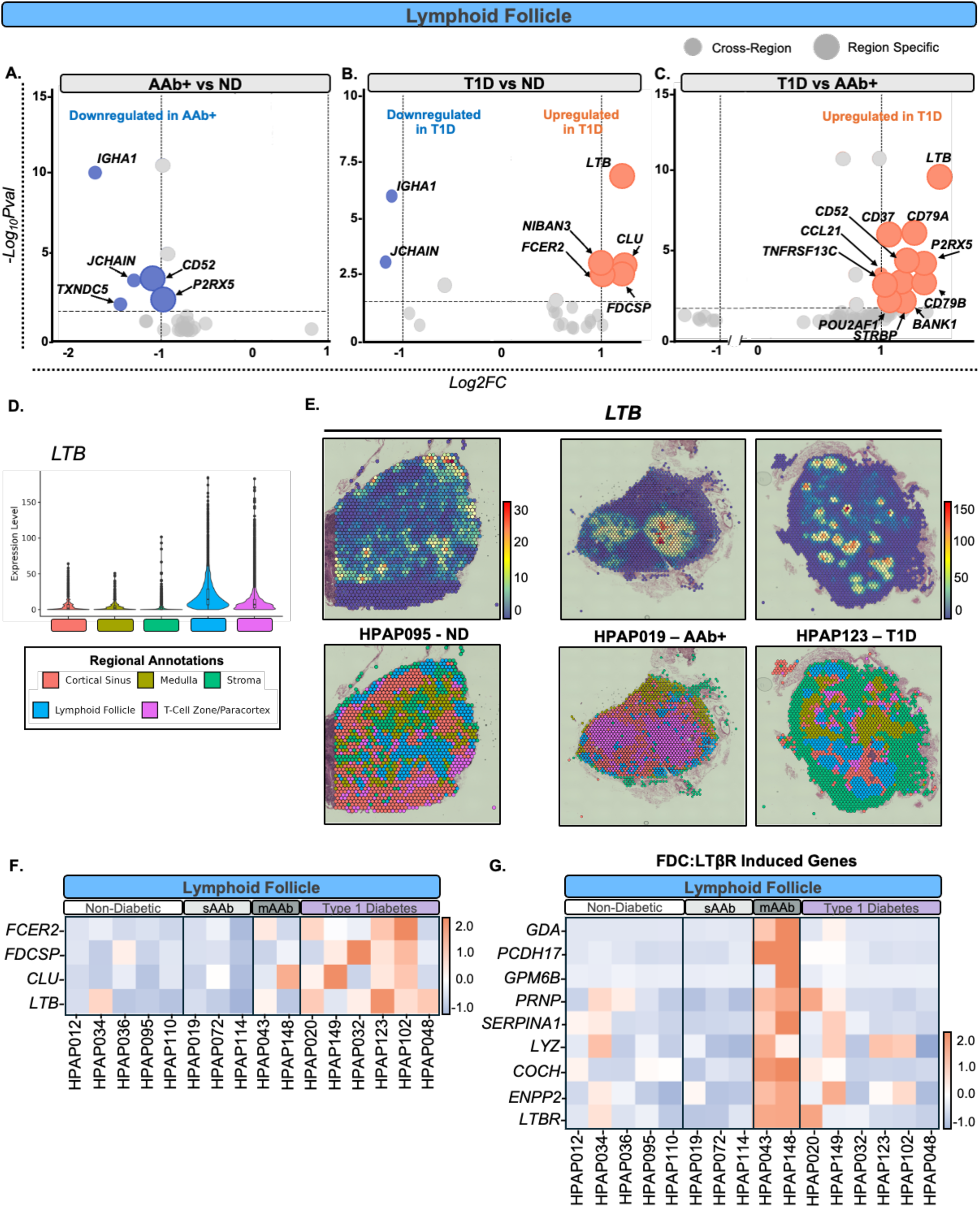
T1D lymphoid follicles exhibit upregulation of lymphotoxin-β and FDC responsiveness genes relative to non-diabetic donors. (A) Volcano plots depicting genes identified as significantly different (adj. p. value < 0.05; Log2FC ≥ |±1|) in the lymphoid follicle between non-diabetic AAB+ donors and ND controls, (B) T1D and ND controls, and (C) T1D and non-diabetic AAb+ sections. Dot size represents whether the DEG was compartment specific (large dot) or detected across multiple pLN compartments (small dot). Upregulated genes are colored in red. Downregulated genes are colored in blue. (D) Violin-Boxplot depicting the normalized expression of *LTB* in each annotated region in donor pLN. Boxplot edges denote the lower and upper quartile with the center dash representing the mean. Dots represent outliers. (E) Spatial profiling of *LTB* expression across representative pLN sections in ND controls, non-diabetic AAB+, and T1D sections (left to right). Corresponding spatially mapped compartment annotations are represented below for each representative section. (F) Heatmap depicting the relative scaled expression of FDC-associated transcripts (*CLU, FDCSP*, and *FCER2*) and *LTB* in the lymphoid follicle across individual donors. (G) Heatmap depicting the relative scaled expression of a previously reported gene set of FDC-specific LTBR inducible genes in the lymphoid follicles across individual donors.

*LTB* is abundantly expressed by T-cells and B-cells^60–62^ and accordingly, the highest regional expression of *LTB* localized to the lymphoid follicles and TCZ (**Fig. 6D**). *LTB* expression across disease status in the pLN displayed the expected spatial expression pattern in ND sections, which is characterized by higher expression in the lymphoid follicles, followed by moderate expression in the adjacent TCZ region (**Fig. 6E**). *LTB* expression was not significantly different in the follicles of AAb+ donors nor in the TCZ of AAb+ and T1D donors relative to ND (**Supp. Fig 6A-B, Supp. Table 4 & 5**). In support of our spatial findings, analysis of the pLN CITE-seq data set (GSE221787) demonstrated significant upregulation of *LTB* among naïve B cell subsets from T1D donors when compared to ND. Interestingly, this analysis revealed that *LTB* was significantly upregulated among naïve and activated effector CD4^+^ T cell subsets and the CD8^+^ naïve #2 T cell subset in AAb+ donors relative to ND (**Supp. Fig. 6C**).

Alongside *LTB*, we demonstrated *CLU* (clusterin)*, FDCSP* (FDC secreted protein), and *FCER2* (Fc epsilon receptor 2) among the top upregulated genes in T1D follicles relative to ND (**Fig. 6B**). Collectively, these genes have previously been described as enriched in mature human follicular dendritic cells (FDCs)^63^, specialized antigen presenting cells (APCs) of stromal lineage that modulate follicular B cell activity and rely on lymphotoxin signals for differentiation^64,65^. Visualizing the expression of these genes revealed a consistent phenotype across individual T1D donors (**Fig. 6F**).

To assess whether LTβ signals were associated with FDC responsiveness in the T1D lymphoid follicles, we measured gene expression of a previously reported FDC-specific LTβ receptor (LT-βR) inducible gene set across donors^66^. Strikingly, the LTβR inducible gene set appears uniformly upregulated in lymphoid follicles of the two mAAb+ donors relative to all other groups, albeit with limited enrichment in T1D donors of shorter disease duration (**Fig. 6G**). Interestingly, one new-onset T1D donor (HPAP020; 4 days duration), exhibited elevated expression of *LTBR* (**Fig. 6G**). Taken together, these results reveal a previously unreported enrichment of FDC-associated transcripts in the lymphoid follicles of AAb+ donors that may be driven in part by LTβ pathway.

## DISCUSSION

Herein, we report an *in situ* transcriptome-wide reference dataset generated from matched pancreas and pLN tissue sections from human organ donors at distinct stages of T1D. By establishing capture spot annotations using the underlying histological features observed in each tissue sections, we avoid the reliance in inferred cell type identities and were able to define gene expression signatures characteristic of established functional tissue compartments in the pancreas and pLN. Intracompartment differential expression testing across disease status revealed that DEGs exhibit compartment-restricted and globally dysregulated expression patterns. Collectively, these results provide evidence of a distinct inflammation-associated signature in the T1D pancreas and enhanced follicular activity in the T1D pLN.

Notably, we identified a previously undescribed increase in *LTB* expression within the pLN lymphoid follicles of T1D donors relative to ND and AAb+ donors. *LTB* encodes LTβ, a tumor necrosis factor superfamily (TNFSF) member involved in lymphoid organogenesis and homeostatic maintenance of secondary lymphatic tissue^67^. A role for LTβ in the pathogenesis of autoimmune diabetes was previously observed when the administration of soluble LTβR-Ig in non-obese diabetic (NOD) mice was found to delay disease onset and prevented the formation of pancreatic tertiary lymphoid structures (TLSs)^68^. Similarly, soluble LTβR-Ig treatment during NOD gestation also conferred complete protection from disease out to one year of age in the offspring. Remarkably, adoptive transfer of bulk splenocytes from gestationally LTβR-Ig treated NODs failed to transfer disease in 12/13 NOD.*scid* recipients^69^. These findings from murine models, taken together with our results in human tissues, implicate the LTβ signaling pathway as a key candidate for therapeutic targeting. Indeed, our data in both AAb+ and T1D donors would suggest potential benefit from agents capable of blocking both LTβ and TNF (e.g., etanercept, a blocking soluble receptor-Ig fusion protein)^70^, versus agents that singularly block TNF or its receptor, which have also demonstrated some efficacy in stage 3 T1D^71^.

In the T1D lymphoid follicles, we also observed increased expression of *CLU, FCER2,* and *FDCSP*, which have been previously characterized as human FDC marker genes^63^. Interestingly, when we measured the expression of a previously reported FDC-specific LTβR inducible gene set across donors^66^, this signature was enriched most strongly in mAAb+ donors, with a weaker signal in recent onset T1D cases. This suggests the LTβR inducible gene set is enriched before *LTB* upregulation and enrichment of FDC-associated maturation genes. Our spatial data did not reveal a change in *LTB* expression before disease onset; however, re-analysis of published pLN CITE-seq data^3^ revealed significant upregulation of *LTB* among several naïve CD4^+^ and CD8^+^ and activated CD4^+^ T cell subsets in AAb+ donors (**Supp. Fig. 7C**).

Functionally, LTβ-mediated signaling involves interactions between lymphocytes and cells of stromal lineage which are critical regulators of immune cell trafficking and maintenance of the lymphoid microenvironment via direct regulation of chemokine production^72^. Elevated LTβ expression may contribute to the pathogenesis of T1D by distinct mechanisms in the pancreas and pLN. Within the pLN, increased LTβ in the lymphoid follicles may increase the activation state of follicular FDCs and augment their antigen presenting capabilities or increase the expression of costimulatory molecules facilitating the activation of autoreactive lymphocytes. Alternatively, increased LTβ expression on the surface of pancreas-infiltrating lymphocytes may promote further immune infiltration or the organization of TLSs by inducing local chemokine production from pancreatic endothelial or stellate cells. Alongside our observations of upregulated *LTA* expression within immune enriched capture spots in the sAAb+ pancreas, this further implicates lymphotoxin signaling as a key inflammatory pathway prior to disease onset.

Analysis of pancreas sections also revealed significant changes in the expression of several *REG* family members before and after disease onset. The *REG* gene family, initially identified in regenerating pancreatic islets in rats^73^, consists of five members in humans (*REG1A, REG1B, REG3A, REG3G*, and *REG4*), each of which encode a small, secreted C-type lectins^73^. Prior research has shown a clear association between *REG* expression and inflammation, as evidenced by multiple reports documenting increased expression of *REG1* and *REG3* orthologs in murine models of acute pancreatitis^74,75^. Overexpression of REG genes has also been reported in pancreata from NOD mice and new-onset human T1D^76,77^. These reports are concordant with our observations, which featured significantly greater expression of *REG1B* and *REG3A* in human T1D across all pancreas compartments. Our analyses also revealed upregulation of *REG3G* in T1D sections relative to ND donors, however, this was only specific to the exocrine compartment. Interestingly, all AAb+ donors in our analyses had significantly lower tissue-wide *REG* expression relative to T1D and ND donors, suggesting their involvement both pre- and post-onset. REG proteins have trophic and protective qualities on islet cells^78,79^. Specifically, REG3B has been shown to protect against streptozotocin-induced β-cell death^80–82^. Hence, reduced REG expression in AAb+ pancreas may signify the loss of protective factors against β-cell death while REG upregulation post-onset may reflect pancreatic inflammation or survival bias of residual cells.

The endocrine compartment featured the majority of the pancreas compartment-specific DEGs. We detected downregulation of *INS* and *GCG* in non-diabetic AAb+ sections relative to ND which implies deficiencies in endocrine function occur prior to disease onset which is consistent with prior findings^23,26,95^. We also detected reduced expression of *CUZD1, TTR,* and *SERPINA3* among AAb+ donors relative to ND. *CUZD1* encodes a zymogen granule-associated transmembrane protein highly expressed in the acinar pancreas with implications in pancreatitis^96^. Due to the expected multicell composition of individual capture spots, it is likely that endocrine compartment-specific downregulation of *CUZD1* may reflect gene expression changes specifically within islet-adjacent acinar cells in AAb+ donors since *CUZD1* expression is unchanged in the acinar compartment. *TTR* encodes transthyretin which, in addition to its more well-known roles in retinol and thyroxine transport, is also expressed in the islets by both α- and β-cells^97^ where it participates in the secretion of glucagon^98^ and insulin^27^, respectively. *SERPINA3* encodes a serine protease inhibitor that has been shown previously to promote insulin production in response to a high-fat diet by negatively regulating the JNK pathway^99^.

The study of established T1D cases was limited by a paucity of insulitic lesions, as confirmed by expert histopathological assessment (**Supp. Table 3**). Insulitis presents with a heterogenous pattern of inflammation between lobules of the same pancreas and across samples and is most prevalent within the first year following T1D onset^100^. Despite this, cell deconvolution was employed to identify capture spots exhibiting enrichment of immune-associated transcripts in order to focus on areas of the pancreas suspected to be involved in some inflammation response. Using this approach, we detected the most immune-enriched capture spots in mAAb+ pancreas relative to ND, sAAb+, and T1D groups. We observed the most abundant enrichment of general T cell markers in mAAb+ donors, whereas T1D donors were predominantly enriched with general B cell markers. The latter finding is consistent with the higher frequency of proliferating B cells in the T1D pancreas as observed by imaging mass cytometry^35^.

Within pLN sections, naïve lymphocyte, myeloid APC, and ILC subsets were predominantly enriched in ND donors and relatively infrequent in T1D donors. Various activated, effector, and central memory T cell subsets exhibited peak enrichment in mAAb+ but were infrequent in T1D donor pLN. Interestingly, several memory B cell phenotypes were uniquely identified as enriched among T1D donors, indicating that humoral immune activity may be persistently augmented in the pLN in established disease. These findings match our observations in donor-matched pancreas sections, indicating a possible common B cell signature across both tissues. Contrary to these observations in tissues, several peripheral blood studies have reported either no significant difference or reduced memory B cell frequencies in T1D compared to controls^101–104^. This points towards differences between systemic circulation and tissue-specific accumulation.

By leveraging the positionally resolved gene expression imparted by ST, we uncovered both global and compartmentally localized changes in gene expression, including both immunomodulatory and functional pancreas genes, from paired pancreas and pLN acquired from a cross-sectional cohort of human donors at distinct stages along the natural progression of T1D. Furthermore, by leveraging complementary publicly available gene expression datasets for *in silico* cell deconvolution, we identified shifts in immune phenotypes and cellular frequencies across both tissues, which can aid in the identification of cellular targets or inflammatory pathways for the development of preventative therapies at elevated risk of disease progression. Lastly, these data represent a valuable resource for the T1D-community providing tissue-specific inflammation signatures that can be linked and integrated with additional molecular readouts (e.g., high parameter proteomics, epigenetics, metabolomics) to create a comprehensive multi-modal description of inflammation and pancreatic dysfunction across the pathogenic stages of T1D.

## LIMITATIONS OF THE STUDY

There are several limitations to this study that should be acknowledged. First, our findings our based on a cross-sectional cohort design. While we aimed to include representative donors across the natural progression of T1D, our characterizations are limited to a single time point. Furthermore, while our study provides spatial context to identified gene expression profiles, our findings are subject to sampling bias caused by profiling individual tissue sections which may not fully capture the cellular heterogeneity present throughout entire tissues. Finally, multi-cell resolution ST inherently lacks single-cell resolution and reflects the aggregate transcriptome of multiple cells at each location. While integration with single cell transcriptome data facilitated the interpretation of these data at the cellular level, our findings require additional validation with single-cell or subcellular resolution ST or the incorporation of orthogonal validation studies to further support these findings.

## RESOURCE AVAILABILITY

### Lead Contact

Requests for further information and resources should be directed to, and will be fulfilled by the lead contact, Todd M. Brusko, PhD.

### Materials Availability

This study did not generate new unique reagents. Donor tissues evaluated in this study were obtained from the HPAP consortium and are available for HPAP investigators.

### Data and Code Availability

FASTQ and image files associated with the data reported herein are deposited at the Gene Expression Omnibus (GEO) repository under accession number GSE296626 and will be made available on the HPAP data portal (https://hpap.pmacs.upenn.edu/). This paper does not report original code. Any additional information required to reanalyze the data reported in this paper is available from the lead contact upon request.

## ACKNOWLEDGEMENTS

This research was supported by grants from the National Institutes of Health (NIH) HIRN-HPAP, P01 AI042288 to TMB, T32 5T32DK108736-08 to MAM-S, R01DK123329 to MCT. This manuscript used publicly available data generated by the Human Pancreas Analysis Program (HPAP-RRID:SCR_016202) database (https://hpap.pmacs.upenn.edu), a Human Islet Research Network (RRID:SCR_014393) consortium (UC4-DK-112217, U01-DK-123594, UC4-DK-112232, and U01-DK-123716). We thank Dr. Vincent Wu (University of Pennsylvania) for providing cell type annotations for the pLN CITE-seq reference data. The University of Florida Interdisciplinary Center for Biotechnology Research (UF | ICBR) provided sequencing support for ST libraries.

## AUTHOR CONTRIBUTIONS

Miguel A. Medina-Serpas: Investigation, formal analysis, data curation, writing- original draft, visualization

Maigan Brusko: Methodology, writing- reviewing and editing, supervision, project administration

Gregory Golden: Case selection. writing- reviewing and editing, analysis and annotation of pLN CITE-seq data

Martha Campbell-Thompson: Investigation, data curation, writing- reviewing and editing

Trevor Rogers: Investigation, writing- reviewing and editing

Eline T. Luning Prak: Writing- reviewing and editing

Chengyang Liu: Tissue sample processing

Klaus Kaestner: Resources, writing- reviewing and editing, funding acquisition

Ali Naji: Resources, writing- reviewing and editing, funding acquisition

Lauren McIntyre: Formal analysis, writing- reviewing and editing

Michael Betts: Conceptualization, resources, writing- reviewing and editing, funding acquisition

Mark A. Atkinson: Resources, writing- reviewing and editing, funding acquisition

Todd M. Brusko: Conceptualization, methodology, resources, writing- reviewing and editing, supervision, funding acquisition

## SUPPLEMENTAL FIGURE LEGENDS

**Figure S1:**
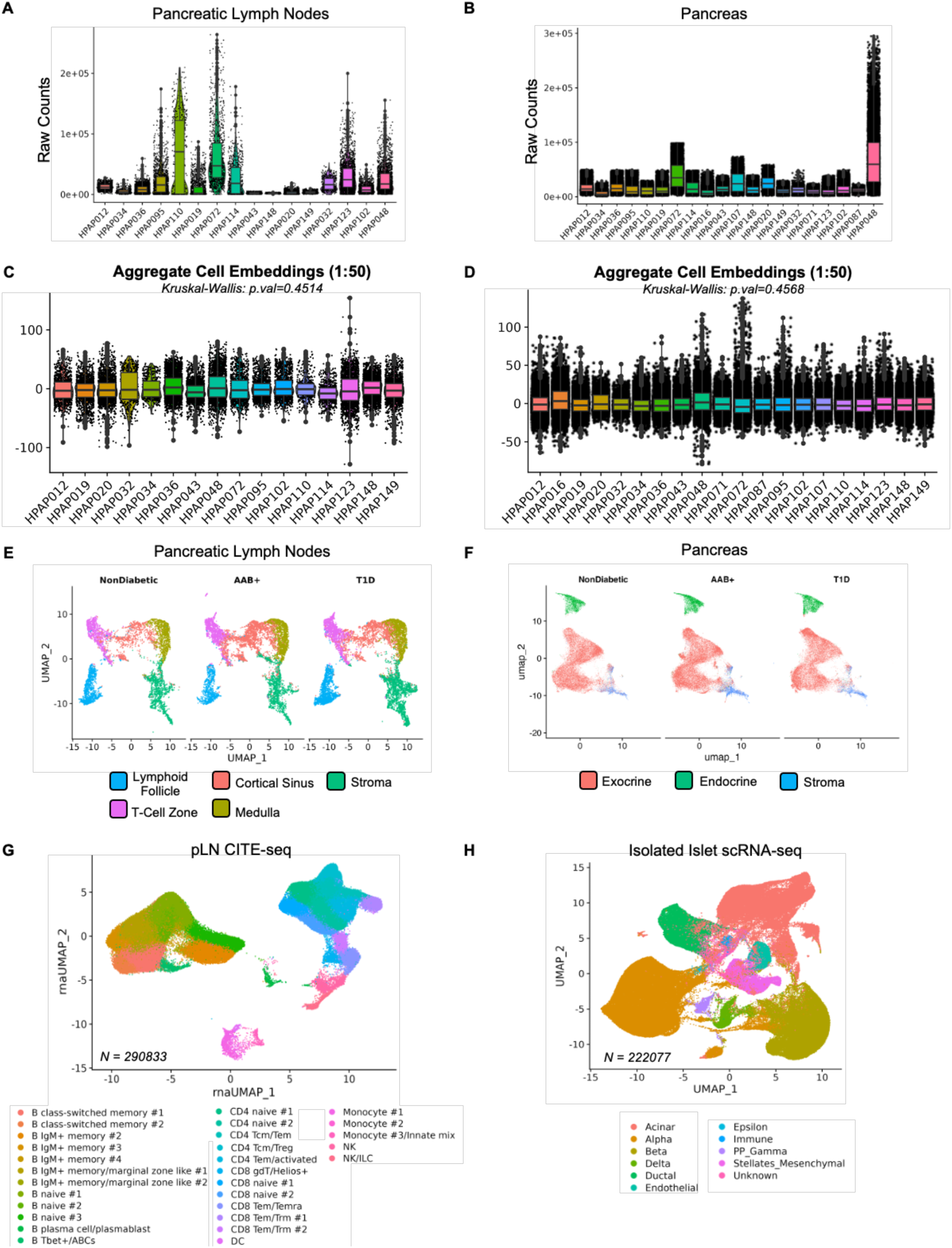
(A) Violin-boxplot depicting the unnormalized count distribution for pLN and (B) pancreas spatial transcriptomic libraries across individual donors. Boxplot edges denote the lower and upper quartile with the center dash representing the mean. Dots represent outliers. (C) Violin-boxplot depicting the aggregate embedding values from the first 50 dimensions for each capture spot following principal component analysis (PCA) in the pLN and (D) the pancreas. Boxplot edges denote the lower and upper quartile with the center dash representing the mean. Dots represent outliers. Significance was calculated by Kruskal-Wallis test. (E) Annotated pLN and (F) pancreas capture spots represented in 2-dimensions as a uniform manifold approximation projection (UMAP) post integration across disease status. (G) UMAP representation of annotated cell subsets identified in the pLN CITE-seq reference data set and (H) isolated islet pancreas scRNA-seq reference data set.

**Figure S2:**
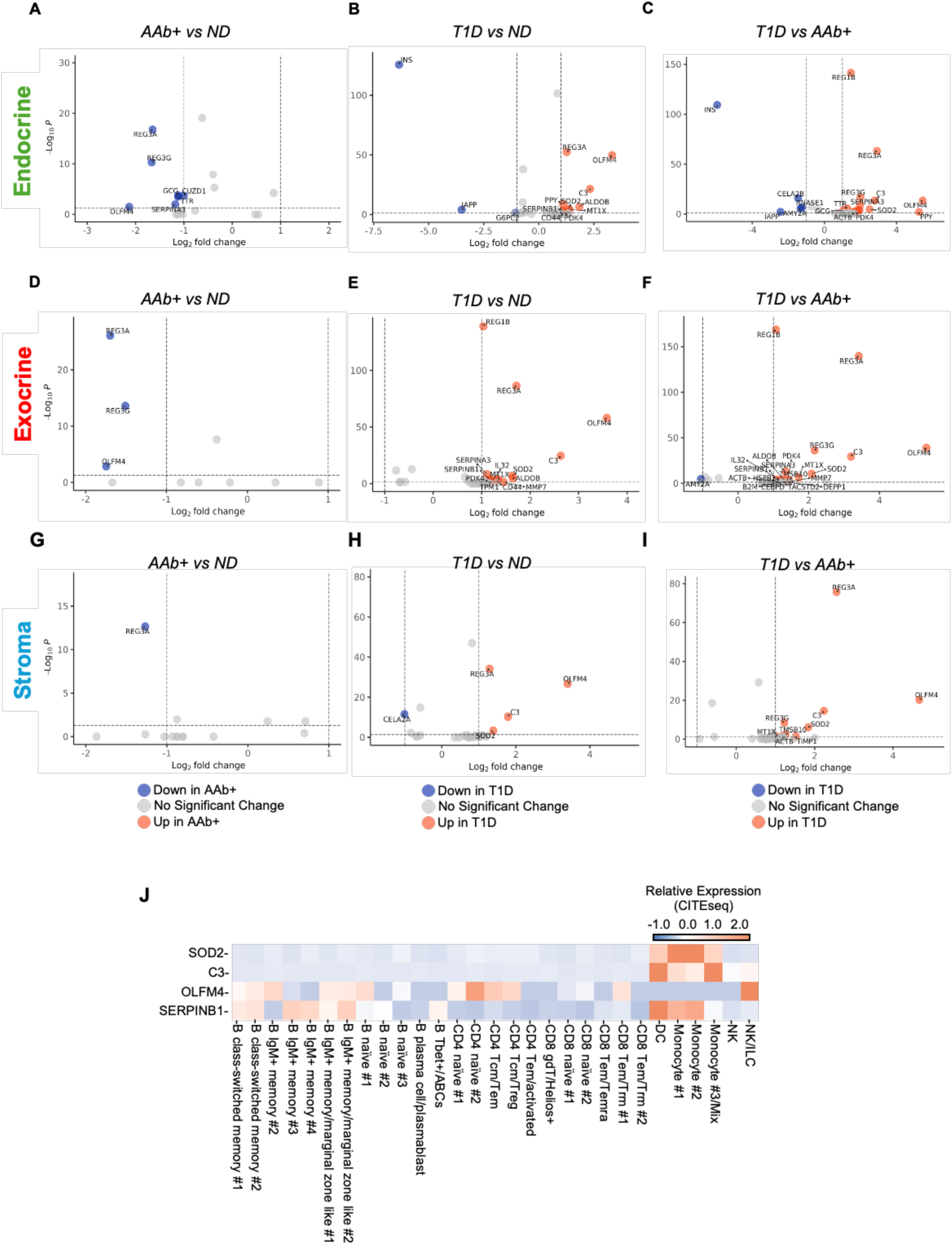
Volcano plots of up- and down-regulated genes identified in the endocrine compartment between (A) AAb+ and ND controls, (B) T1D and ND controls, and (C) T1D and AAb+ donors. Volcano plots of up- and downregulated genes identified in the exocrine compartment between (D) AAb+ and ND controls, (E) T1D and ND controls, and (F) T1D and AAb+ donors. Volcano plots of up- and downregulated genes identified in the stromal compartment between (G) AAb+ and ND controls, (H) T1D and ND controls, and (I) T1D and AAb+ donors. Red points indicate upregulated genes. Blue points indicate downregulated genes. Genes with an adjusted p value < 0.05 (Benjamini-Hochberg correction) are considered significant and genes with an absolute Log_2_FC > 1 are displayed. (J) Heatmap depicting the relative scaled expression of *SOD2, OLFM4, C3,* and *SERPINB1* across annotated immune subsets identified in the pLN CITE-seq reference data set.

**Figure S3:**
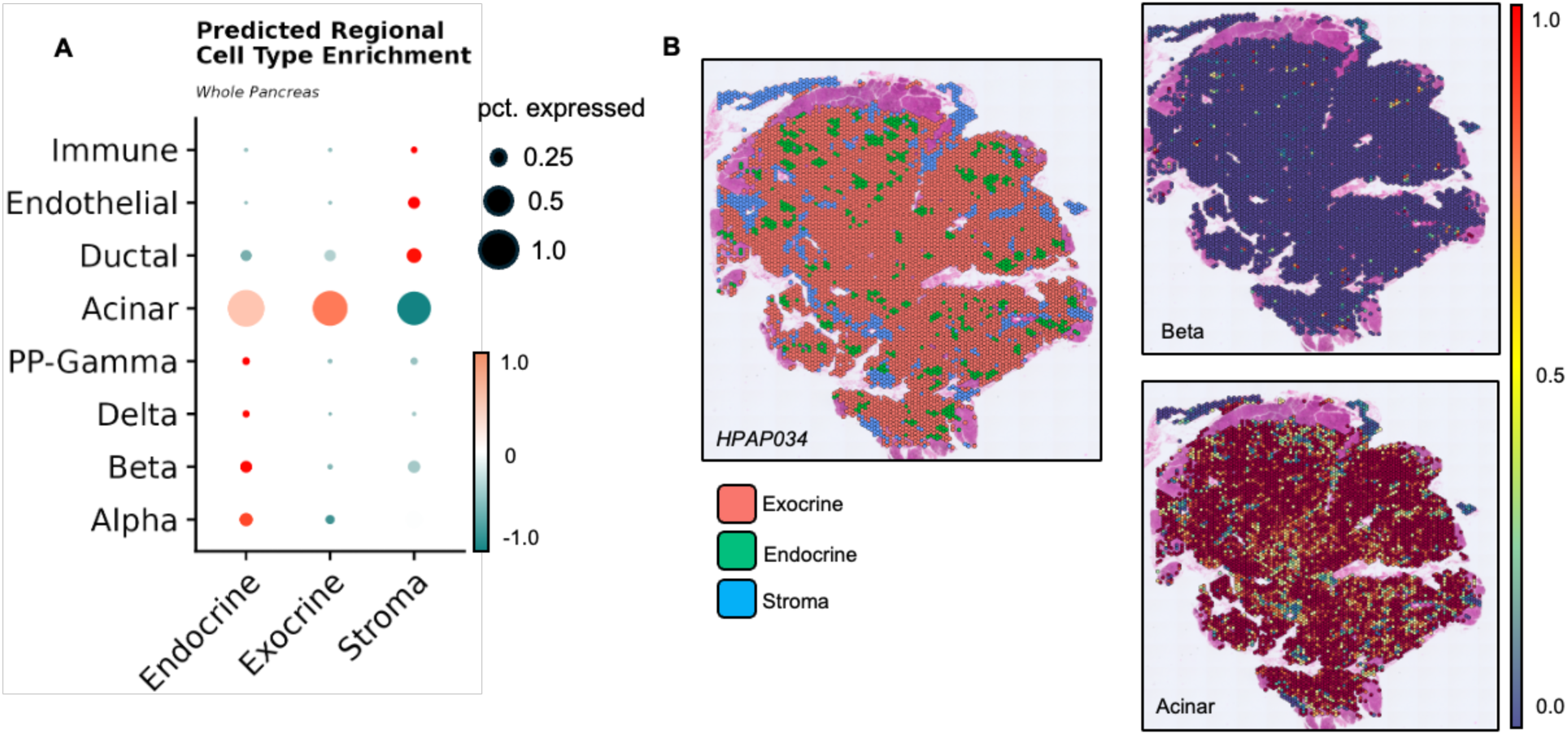
(A) Dot plot depicting the relative scaled enrichment of each annotated cell subset represented in the pancreas scRNA-seq reference data set by cell deconvolution across each annotated regional compartment defined in the pancreas ST data. Dot size is representative of the predicted cell type frequency within each annotated regional compartment. (B) Spatial projection of annotated pancreas capture spots across a representative tissue section defined by ST (left). The predicted cell type frequency of β-cells (top) and acinar cells (bottom) was calculated by cell deconvolution and spatially projected onto the corresponding representative section. Scale is represented as an absolute scale from 0.0 (blue) to 1.0 (red) where the value represents the predicted cell-type proportion contained within each individual capture spot.

**Figure S4:**
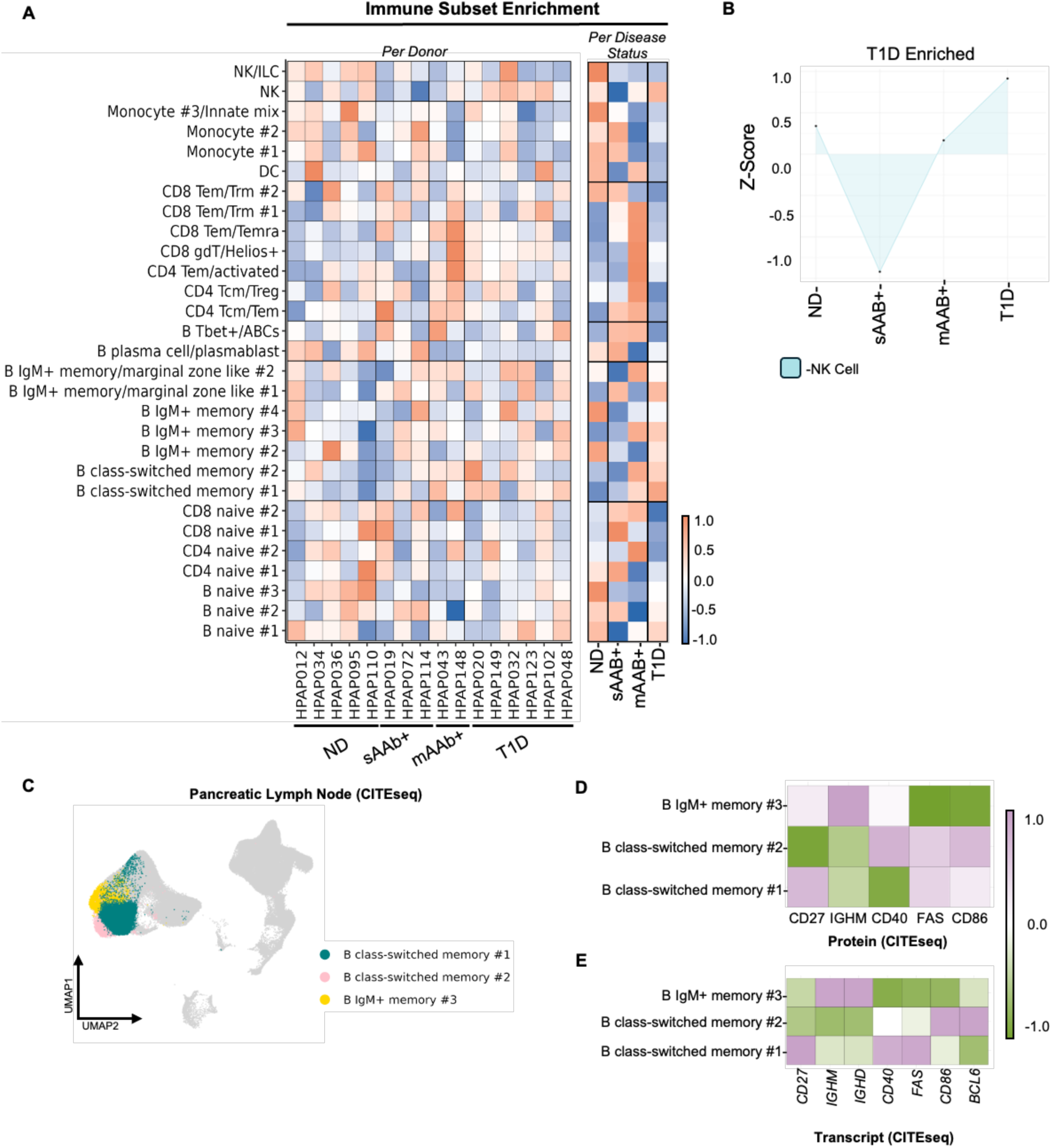
(A) Heatmap depicting the relative scaled enrichment of each annotated cell subset represented in the pLN CITE-seq reference across individual donors upon cell deconvolution of the pLN ST data set. (left). Relative cell type enrichment is also represented across disease status (right). (B) Cell type enrichment module representative of T1D-enriched only cell types. Each point is representative of the relative scaled expression at each disease classification. (C) UMAP projection of three annotated B-cell subsets defined in the pLN CITE-seq reference that were identified as enriched in the mAAb+/T1D module. (D) Heatmap depicting the relative scaled protein marker and (E) corresponding gene expression of memory, class-switch, and costimulatory markers from mAAb+/T1D enriched module B-cell subsets in the pLN CITE-seq reference.

**Figure S5:**
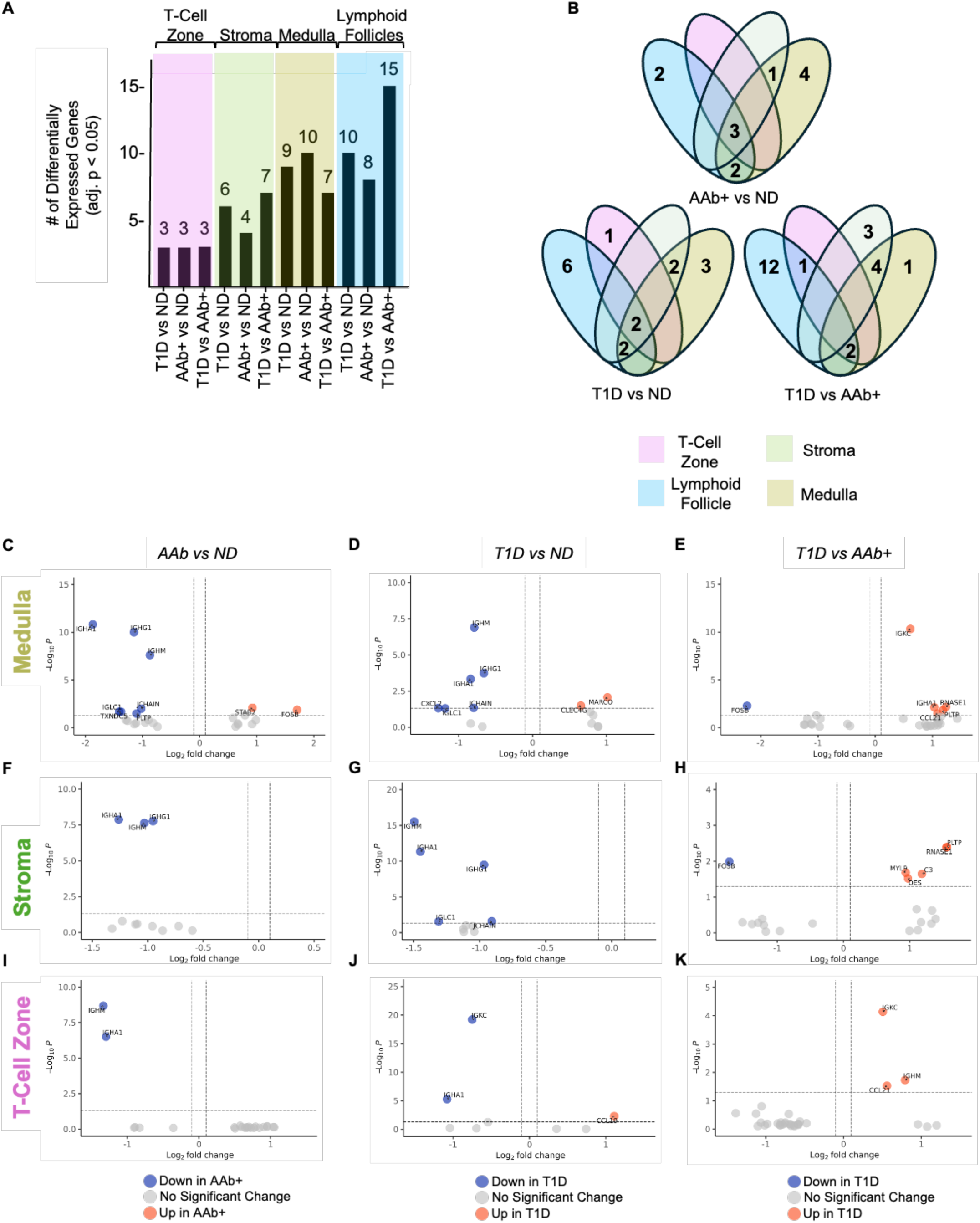
(A) Bar plot displaying the number of significant differentially expressed genes within each annotated pLN compartment across each pairwise comparison. (B) Venn diagram depicting the number of significant differentially expressed genes that are unique to or shared across each annotated pLN compartment. (C) Volcano plots of up- and down-regulated genes identified in the medulla compartment between AAb+ and ND controls, (D) T1D and ND controls, and (E) T1D and AAb+ donors. Volcano plots of up- and downregulated genes identified in the stromal compartment between (F) AAb+ and ND controls, (G) T1D and ND controls, and (H) T1D and AAb+ donors. Volcano plots of up- and downregulated genes identified in the T-cell zone compartment between (I) AAb+ and ND controls, (J) T1D and ND controls, and (K) T1D and AAb+ donors. Red points indicate upregulated genes. Blue points indicate downregulated genes. Genes with an adjusted p value < 0.05 (Benjamini-Hochberg correction) are considered significant and genes with an absolute Log_2_FC > 1 are displayed.

**Figure S6:**
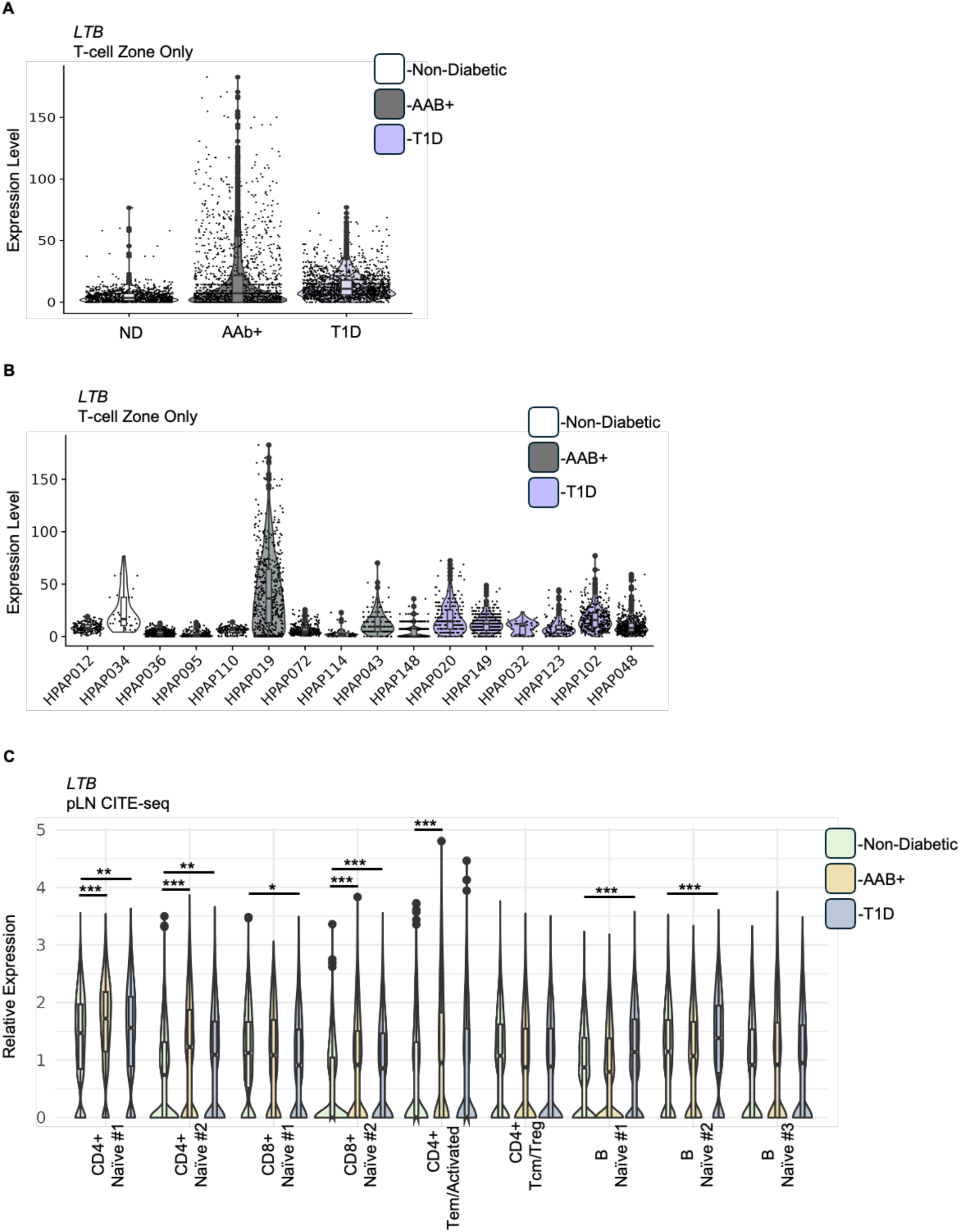
(A) Violin-boxplot depicting the normalized expression of *LTB* in the T-cell zone across disease status. (B) Violin-boxplot depicting the normalized expression of *LTB* in the T-cell zone across individual donors. (C) Violin boxplot depicting the normalized expression of *LTB* across a variety of annotated T-cell and B-cell subsets described in the pLN CITE-seq reference data set. Boxplot edges denote the lower and upper quartile with the center dash representing the mean. Dots represent outliers. Significance was determined by students t-test across three pairwise comparisons; AAb+ vs ND, T1D vs ND, and T1D vs AAb+. *p value < 0.05, **p value < 0.005, ***p value < 0.0005.

## SUPPLEMENTAL INFORMATION

**Document S1.** Figures S1-S6 and Table S2-5, S8-10

**Table S1.** Donor summary characteristics. Related to Figure 1.

**Table S2.** Histopathology notes of pancreatic lymph node sections. Related to STARMethods and Figure 1.

**Table S3.** Histopathology notes of pancreas sections. Related to STARMethods and Figure 1.

**Table S4.** Differential expression table corresponding to the endocrine compartment in the pancreas. Related to Figure 2.

**Table S5.** Differential expression table corresponding to the exocrine compartment in the pancreas. Related to Figure 2.

**Table S6.** Differential expression table corresponding to the stromal compartment in the pancreas. Related to Figure 2.

**Table S7.** Differential expression table corresponding to the lymphoid follicle compartment in the pLN. Related to Figure 6.

**Table S8.** Differential expression table corresponding to the T-cell zone (TCZ) compartment in the pLN. Related to Figure 6.

**Table S9.** Differential expression table corresponding to the stromal compartment in the pLN. Related to Figure 6.

**Table S10.** Differential expression table corresponding to the medulla compartment in the pLN. Related to Figure 6.

## Methods

### EXPERIMENTAL MODEL AND STUDY PARTICIPANT DETAILS

#### Human Organ Donor Tissues

Pancreas and pLN FFPE sections were obtained from the HPAP consortium (RRID:SCR_016202; https://hpap.pmacs.upenn.edu), part of the Human Islet Research Network (https://hirnetwork.org/), with approval from the University of Florida Institutional Review Board (IRB # 201600029) and the United Network for Organ Sharing (UNOS). A legal representative for each donor provided informed consent prior to organ retrieval. Donor summary characteristics are outlined in **Supplemental Table 1**.

### METHOD DETAILS

#### Spatial Transcriptomics

Spatial transcriptomics of pancreatic and pLN tissues were performed using the Visium CytAssist Spatial Gene Expression for FFPE kit (10x Genomics) according to the manufacturer’s protocol. Briefly, FFPE tissue blocks from pancreas and pLN tissue for each donor were cut to a thickness of 5µm and mounted onto positively charged microscope slides. Tissue sections were deparaffinized and rehydrated according to Visium CytAssist Tissue Preparation Guide Tissue sections were H&E stained and imaged using either a BZ-X700 (Keyence) or PhenoImager Slide Scanner (Akoya Biosciences) for histological assessment and gene expression underlay. Whole transcriptome probe hybridization was performed for approximately 18 hours at 50°C for both tissue types. Probe hybridized tissue sections were transferred onto a 6.5mm x 6.5mm (pLN) or 11mm x 11mm (pancreas) Visium CytAssist Gene Expression slide using the Visium CytAssist instrument with transfer conditions set to 37°C for 30 minutes for both tissues. Spatially barcoded cDNA libraries were analyzed by Agilent High Sensitivity DNA kit on the 2100 Bioanalyzer (Agilent) to estimate library concentration and quality. Libraries were pooled to equimolar concentrations and sequenced on an S4 Illumina flow cell using the NovaSeq 6000 (Illumina) with a target depth of 25,000 reads per tissue-covered capture spot. Arrangement of donor sections on the Visium CytAssist Spatial Gene Expression array is provided in **Supp. Table 1**. FASTQ and image files are deposited at the Gene Expression Omnibus (GEO) repository under accession number GSE296626.

#### Histopathology review

Paraffin sections were stained by H&E and whole slide scans reviewed for pathological findings. Tissue sections were evaluated using a semi-quantitative scoring system (0= none or absent, 1= few or mild, 2= several or moderate, 3= many or severe) as previously reported^105^. pLN was evaluated for adequacy of tissue compartments (follicles, medulla, cortex) and the presence of fat, fibrosis or other significant findings (**Supp. Table 2**). Pancreas was evaluated for relative numbers of islets, acute or chronic inflammation (endocrine and exocrine compartments), acinar atrophy, fat, fibrosis and other features (**Supp. Table 3**).

#### Immunofluorescence

Immunofluorescence staining was performed according to the PhenoCycler-Fusion user guide (Akoya Biosciences). Deparaffinization and rehydration of FFPE tissue was performed as described with exception for our choice of solvent, xylene was used instead of HistoChoice Clearing Agent. Antigen retrieval was performed by immersing sections in 1x AR9 antigen retrieval buffer (Akoya Biosciences, PN#AR6001K) and incubating for 20 minutes under high pressure in a pressure cooker. Primary antibody staining using oligonucleotide conjugated anti-Hu CD20 (AKYP0049-BX064), anti-Hu CD3 (AKYP0027-BX015), and anti-Hu Ki67 (AKYP0052-BX047) was performed for 3 hours at room temperature. A complementary oligonucleotide conjugated fluorescence reporter plate corresponding to barcodes BX064, BX015, and BX047 were prepared for protein detection. Automated detection and imaging were performed at 40x magnification using the PhenoCycler-Fusion System (Akoya Biosciences).

### QUANTIFICATION AND STATISTICAL ANALYSIS

#### QC, Data Integration, and Normalization

Initial data quality was assessed on a per-tissue and per-donor basis prior to data integration and downstream analysis. Summary count information on each donor and tissue is outlined in **Supp. Table** 1. We evaluated the total number of reads, detected genes, and percentage of reads mapping to mitochondrial genes per sample and per capture spot. Capture spots with mitochondrial content ≥ 10% were removed. Individual donor datasets were integrated by respective tissue (pancreas n=20; pLN n=16) by anchor based canonical correlation analysis (CCA) integration procedure implemented in Seurat to create a unified data set. After integration, unnormalized count data for each donor was normalized by upper quartile normalization^20^. Lastly, expression values for each gene were standardized across all capture spots (z-score transformation) by implementing the *ScaleData* function in *Seurat*.

#### Feature Selection & Dimensionality Reduction

Prior to dimensionality reduction, a total of 1000 highly variable genes detected across all donors were identified for downstream analysis using the *FindVariableFeatures* function in Seurat using the *vst* method (selection.method = ‘vst’). We then performed principal component analysis (PCA) on the top 1000 highly variable genes using the *RunPCA* function to visualize the grouping of each annotated compartment along the top 3 dimensions for each respective tissue. Lastly, cells were clustered and projected in 2-dimensions by uniform manifold approximation projection (UMAP) using the *RunUMAP* function to qualitatively assess the performance of our integration strategy (**Supp. Fig. 1B & 1D**).

#### Tissue Compartment Annotations

Tissue compartment annotations were based on H&E stained tissue sections generated during the Visium assay were used to demarcate major functional regions of the pancreas and pLN. For the pancreas, we identified 3 major tissue compartments: the exocrine, endocrine, and stromal compartment. For the pLN, we identified 5 major tissue compartments: the lymphoid follicles, T-cell zone (TCZ), indistinct region, medulla, and stromal compartment. Upon assigning capture spots into histologically defined compartment level annotations, we identified the top 20 compartment marker genes using *FindAllMarkers* (test.use = ‘wilcox’, min.pct = 0.25). We confirmed tissue compartment annotations are transcriptionally defined by marker genes associated with the predominant cell type known to localize to these regions (**Fig. 2A & 4A**).

#### Differential Expression (DE) Testing

We performed three pairwise comparisons: AAb+ vs ND, T1D vs ND, & T1D vs AAb+. Genes considered for DE testing were detecting in ≥ 25% of capture spots per annotated compartment across all donors. DE genes were identified using the likelihood ratio test on a generalized linear model using the *FindMarker* function in Seurat (test.use = poisson). Nominal p-values were adjusted using the Benjamini-Hochberg procedure for multiple test correction^106^. Genes were reported as significantly DE if the absolute log2FC difference is ≥ 0.25 and has a false discovery rate (FDR) < 0.05. DE analysis results are visualized using *EnhancedVolcano*^107^.

#### Capture Spot Cell Deconvolution

Cell deconvolution of multi-cell resolution spatial transcriptomics in the Visium gene expression assay enables inference of constituent cell type proportions contained within individual capture spots^108^. For this approach, we integrated our individual spatial gene expression assays from pancreas and pLN with a single-cell transcriptomic reference data. The isolated islet pancreas scRNA-seq data were obtained from the the PancDB database^18^. Only samples from T1D individuals were included. The pLN CITE-seq data were acquired from HPAP (GSE221787). These single-cell reference data contain 40% and 50% donor overlap with our pancreas and pLN spatial gene expression cohort, respectively. Donor representation in each respective single cell reference is outlined in **Supp. Table 1**. Deconvolution was implemented using the anchor-based canonical correlation analysis (CCA) method in the *FindTransferAnchors* function in Seurat with parameters set to default. The predicted cell-type proportion at each capture spot position was calculated for each cell type annotation represented in the reference dataset.

#### Statistical Analysis

No statistical test was performed to predetermine sample size. We performed three pairwise comparisons: AAb+ vs ND, T1D vs ND, & T1D vs AAb+. We employed the likelihood ratio test within Seurat. Genes with an absolute log_2_FC > 0.25 and FDR < 0.05 using the Benjamini-Hochberg test correction were reported as significant (Supp. Table 4-10). We used the unpaired students T-test to calculate the significance of cell deconvolution values and CITE-seq RNA expression data between ND controls and T1D or AAb+ donors. We used the Kruskal-Wallis test to calculate the significance of aggregate cell embedding values across individual donors. Significance values are denoted as: *p < 0.05, **p < 0.005, ***p < 0.0005, and ****p < 0.00005. All statistical analyses were performed using R version 4.2.3. Seurat version 4.3.0 was utilized for these analysis.

