## Supplemental Table 1 for "Spatial transcriptomics from pancreas and local draining lymph node tissue reveals a lymphotoxin-β signature in human type 1 diabetes"

Summary Donor Characteristics

| **Disease Status Classification** | | | | | |
| --- | --- | --- | --- | --- | --- |
|  |  | Autoantibody Positive (AAb+) | |  |  |
|  | Non-Diabetic (ND) | Single AAb+ | Multiple AAb+  (≥ 2) | T1D | TOTALS |
| **Total Donors (n)** | 5 | 3 | 4 | 8 | 20 |
| **Sex, n (%)** |  |  |  |  |  |
| Male | 2 (40) | 2 (66.66) | 4 (100) | 5 (62.5) | 13 |
| Female | 3 (60) | 1 (33.33) | 0 | 3 (37.5) | 7 |
| **Age (yr)** |  |  |  |  |  |
| Mean +/- (SD) | 21.6 (5.97) | 22.5 (10.61) | 16.75 (9.60) | 16.5 (6.40) |  |
| Median (Q1-Q3 | 23 | 22.5 | 15 | 14.5 |  |
| **Ethnicity, n (%)** |  |  |  |  |  |
| Non-Hispanic | 5 (100) | 2 (66.67) | 3 (75) | 7 (87.5) | 17 |
| Hispanic | 0 | 1 (33.33) | 1 (25) | 1 (12.5) | 3 |
| **Race, n (%)** |  |  |  |  |  |
| Caucasian | 4 (80) | 3 (100) | 3 (75) | 8 (100) | 18 |
| African American | 1 (20) | 0 | 1 (25) | 0 | 2 |
| Asian | 0 | 0 | 0 | 0 |  |
| Multi | 0 | 0 | 0 | 0 |  |
| **Diagnosis Age (yr)** |  |  |  |  |  |
| Mean +/- (SD) |  |  |  | 12.31 (5.65) |  |
| Median (Q1-Q3 |  |  |  | 10.5 |  |
| **Disease Duration (yr)** |  |  |  |  |  |
| Mean +/- (SD) |  |  |  | 4.19 (2.85) |  |
| Median (Q1-Q3 |  |  |  | 3 |  |
| **HLA Haplotype, n (%)** |  |  |  |  |  |
| DR3/X | 0 | 1 (33.33) | 0 | 1 (12.5) | 2 |
| DR4/X | 1 (20) | 1 (33.33) | 1 (25) | 4 (50) | 7 |
| DR3/4 | 0 | 0 | 1 (25) | 3 (37.5) | 4 |
| DRX/X | 4 (80) | 1 (33.33) | 2 (50) | 0 | 7 |

|  | **Per Donor Summary Characteristics** | | | | | | | | | |
| --- | --- | --- | --- | --- | --- | --- | --- | --- | --- | --- |
|  | **Donor ID** | **Sex** | **Age** | **Race** | **Hispanic/**  **Latino** | **Diagnosis**  **Age (Yrs)** | **Disease**  **Duration (Yrs)** | **# of AABs** | **HLA**  **Diplotype** | **GRS2** |
| **Non-Diabetic** | HPAP012 | Female | 18 | Caucasian | N |  |  | 0 | DR4/X | 11.838 |
|  | HPAP034 | Male | 13 | Caucasian | N |  |  |  | DRX/X | 9.331 |
|  | HPAP036 | Female | 23 | Caucasian | N |  |  |  | DRX/X | 8.804 |
|  | HPAP095 | Female | 23 | AA | N |  |  |  | DRX/X | 9.970 |
|  | HPAP110 | Male | 31 | Caucasian | N |  |  |  | DRX/X | 10.980 |
|  |  | Mean | 21.6 |  |  |  |  |  |  | 10.1846 |
|  |  | Std.Dev | 5.99 |  |  |  |  |  |  | 1.23 |
|  |  | Median | 23 |  |  |  |  |  |  | 9.97 |
| **Single AAb+** | HPAP019 | Male | 22 | Caucasian | N |  |  | 1 | DR4/X | 11.329 |
|  | HPAP072 | Male | 19 | Caucasian | Y |  |  | 1 | DRX/X | 9.157 |
|  | HPAP114 | Female | 21 | Caucasian | N |  |  | 1 | DR3/X | 9.720 |
|  |  | Mean | 20.67 |  |  |  |  |  |  | 10.10 |
|  |  | Std.Dev | 1.53 |  |  |  |  |  |  | 1.13 |
|  |  | Median | 21 |  |  |  |  |  |  | 9.72 |
| **Multiple AAb+** | HPAP107 | Male | 15 | Caucasian | N |  |  | 3 | DR3/4 | 14.118 |
|  | HPAP016 | Male | 30 | Caucasian | N |  |  | 3 | DR4/X | 9.144 |
|  | HPAP043 | Male | 15 | Caucasian | Y |  |  | 2 | DRX/X | 10.990 |
|  | HPAP148 | Male | 7 | AA | N |  |  | 2 | DRX/X | 9.049 |
|  |  | Mean | 16.75 |  |  |  |  |  |  | 10.83 |
|  |  | Std.Dev | 9.60 |  |  |  |  |  |  | 2.37 |
|  |  | Median | 15 |  |  |  |  |  |  | 10.10 |
| **T1D** | HPAP020 | Male | 14 | Caucasian | N | 14 | 0.01 | 4 | DR3/X | 12.770 |
|  | HPAP032 | Female | 10 | Caucasian | N | 7 | 3 | 2 | DR3/4 | 12.968 |
|  | HPAP048 | Male | 27 | Caucasian | Y | 19 | 8 | 0 | DR3/4 | 12.828 |
|  | HPAP087 | Female | 15 | Caucasian | N | 7 | 8 | 1 | DR3/4 | 15.548 |
|  | HPAP102 | Male | 18 | Caucasian | N | 12 | 6 | 1 | DR4/X | 12.742 |
|  | HPAP123 | Male | 25 | Caucasian | N | 22 | 3 | 3 | DR4/X | 11.609 |
|  | HPAP071 | Female | 12 | Caucasian | N | 9 | 3 | 2 | DR4/X | 11.414 |
|  | HPAP149 | Male | 11 | Caucasian | N | 8.5 | 2.5 | 2 | DR4/X | 15.484 |
|  |  | Mean | 16.5 |  |  | 12.31 | 4.19 |  |  | 13.17 |
|  |  | Std.Dev | 6.40 |  |  | 5.65 | 2.85 |  |  | 1.56 |
|  |  | Median | 14.5 |  |  | 10.5 | 3 |  |  | 12.80 |

AA = African American

|  |  |  | Tissue Region of Isolation | |  | Donor Represented in Single Cell Reference Data | |
| --- | --- | --- | --- | --- | --- | --- | --- |
|  | Donor ID |  | Pancreas Region | pLN Region (Relative to Pancreas) |  | Pancreas Islet scRNA-seq | pLN  CITE-seq |
| **Non-Diabetic** | HPAP012 |  | Head | Head |  | No | Yes |
|  | HPAP034 |  | Head | Head |  | Yes | No |
|  | HPAP036 |  | Body | Head |  | Yes | No |
|  | HPAP095 |  | Head | Head |  | No | Yes |
|  | HPAP110 |  | Head | Head |  | No | Yes |
|  |  | #Pan-H | 4/5 | 5/5 |  |  |  |
|  |  | %Pan-H | 80% | 100% |  |  |  |
| **Single AAb+** | HPAP019 |  | Head | Head |  | No | No |
|  | HPAP072 |  | Head | Head |  | Yes | No |
|  | HPAP114 |  | Head | Head |  | No | Yes |
|  |  | #Pan-H | 3/3 | 3/3 |  |  |  |
|  |  | %Pan-H | 100% | 100% |  |  |  |
| **Multiple AAb+** | HPAP107 |  | Head | NA |  | Yes | - |
|  | HPAP016 |  | Head | NA |  | No | - |
|  | HPAP043 |  | Head | Head |  | Yes | Yes |
|  | HPAP148 |  | Head | Head |  | No | No |
|  |  | #Pan-H | 4/4 | 2/2 |  |  |  |
|  |  | %Pan-H | 100% | 100% |  |  |  |
| **T1D** | HPAP020 |  | Head | Head |  | No | Yes |
|  | HPAP032 |  | Head | Head |  | Yes | No |
|  | HPAP048 |  | Head | Head |  | No | Yes |
|  | HPAP087 |  | Tail | NA |  | Yes | - |
|  | HPAP102 |  | Head | Head |  | No | Yes |
|  | HPAP123 |  | Head | Head |  | No | No |
|  | HPAP071 |  | Head | NA |  | Yes | - |
|  | HPAP149 |  | Head | Head |  | No | No |
|  |  | #Pan-H | 7/8 | 6/6 | # in ref | 8/20 | 8/16 |
|  |  | %Pan-H | 87.50% | 100% | % in ref | 40% | 50% |

Pan-H = Pancreas head region

| **Visium Gene Expression Array Pairing of Donor Sections for Processing** | | | | | | | |
| --- | --- | --- | --- | --- | --- | --- | --- |
| pLN (6.5mm x 6.5mm) | | | | Pancreas (11.5mm x 11.5mm) | | | |
| Donor ID | pLN  Visium SN# | Visium Capture Area Position | Library  Construction | Donor ID | Pancreas  Visium SN# | Visium Capture Area Position | Library Construction |
| HPAP012 | V42Y23-322 | A | Batch 1 | HPAP012 | V53M13-019 | A | Batch 1 |
| HPAP020 |  | B | Batch 1 | HPAP072 |  | B | Batch 1 |
| HPAP036 | V42Y23-299 | A | Batch 1 | HPAP087 | V52Y19-315 | A | Batch 1 |
| HPAP034 |  | B | Batch 1 | HPAP123 |  | B | Batch 1 |
| HPAP032 | V42Y23-318 | A | Batch 1 | HPAP016 | V53M20-099 | A | Batch 1 |
| HPAP072 |  | B | Batch 1 | HPAP020 |  | B | Batch 1 |
| HPAP019 | V42Y23-300 | A | Batch 1 | HPAP048 | V53M13-055 | A | Batch 1 |
| HPAP123 |  | B | Batch 1 | HPAP110 |  | B | Batch 1 |
| HPAP095 | V42Y23-347 | A | Batch 1 | HPAP019 | V53M13-023 | A | Batch 1 |
| HPAP102 |  | B | Batch 1 | HPAP034 |  | B | Batch 1 |
| HPAP048 | V42Y93-332 | A | Batch 1 | HPAP102 | V53M20-062 | A | Batch 1 |
| ~~HPAP087~~ |  | B | Batch 1 | HPAP114 |  | B | Batch 1 |
| HPAP110 | V42Y23-394 | A | Batch 1 | HPAP095 | V52Y17-391 | A | Batch 1 |
| HPAP114 |  | B | Batch 1 | HPAP107 |  | B | Batch 1 |
| HPAP043 | V43F23-372 | A | Batch 2 | HPAP032 | V52Y17-318 | A | Batch 1 |
| ~~HPAP071~~ |  | B | Batch 2 | HPAP036 |  | B | Batch 1 |
| HPAP148 | V43F16-130 | A | Batch 2 | HPAP043 | V53U05-315 | A | Batch 2 |
| HPAP149 |  | B | Batch 2 | HPAP071 |  | B | Batch 2 |
| ~~Strikethrough~~: failed to amplify | | | | HPAP148 | V53U05-294 | A | Batch 2 |
|  |  |  |  | HPAP149 |  | B | Batch 2 |

SN# = Serial number

| **Summary Count Table – Pancreas** | | | | | |
| --- | --- | --- | --- | --- | --- |
| Donor ID | Total Counts | Total Capture Spots | Count Q3 | Total Normalized Counts | Normalized Counts per Cell |
| HPAP012 | 139474475 | 7992 | 21360.25 | 117721870 | 14729.96 |
| HPAP016 | 48916377 | 5681 | 11365.00 | 77598547 | 13659.31 |
| HPAP019 | 72924749 | 5430 | 17082.00 | 76967073 | 14174.42 |
| HPAP020 | 37945021 | 1480 | 34036.75 | 20099047 | 13580.44 |
| HPAP032 | 14982440 | 1166 | 16853.25 | 16027568 | 13745.77 |
| HPAP034 | 41706204 | 6475 | 8761.50 | 85820481 | 13254.13 |
| HPAP036 | 120714115 | 6938 | 22616.00 | 96230116 | 13870.01 |
| HPAP043 | 97953133 | 6483 | 18501.00 | 95453504 | 14723.66 |
| HPAP048 | 750846455 | 10346 | 99246.75 | 136396614 | 13183.51 |
| HPAP071 | 45199865 | 4648 | 12376.25 | 65844121 | 14166.12 |
| HPAP072 | 312228460 | 7741 | 57428.00 | 98020642 | 12662.53 |
| HPAP087 | 39758926 | 3296 | 14330.00 | 50021557 | 15176.44 |
| HPAP095 | 65808279 | 4805 | 18959.00 | 62579754 | 13023.88 |
| HPAP102 | 72530014 | 5084 | 18673.75 | 70025299 | 13773.66 |
| HPAP107 | 92027032 | 3404 | 41293.00 | 40179796 | 11803.70 |
| HPAP110 | 86818384 | 7780 | 16882.25 | 92715025 | 11917.10 |
| HPAP114 | 81738649 | 4797 | 25339.00 | 58157634 | 12123.75 |
| HPAP123 | 44759184 | 4591 | 11859.00 | 68046037 | 14821.62 |
| HPAP148 | 143481760 | 10816 | 17556.75 | 147340221 | 13622.43 |
| HPAP149 | 48180222 | 4403 | 13249.50 | 65559866 | 14889.82 |

| **Summary Count Table – pLN** | | | | | |
| --- | --- | --- | --- | --- | --- |
| Donor ID | Total Counts | Total Capture Spots | Count Q3 | Total Normalized Counts | Normalized Counts per Cell |
| HPAP012 | 14350078 | 1123 | 15994.00 | 13001562 | 11577.526 |
| HPAP019 | 15161472 | 1637 | 11977.00 | 18343900 | 11205.803 |
| HPAP020 | 6621537 | 2306 | 4009.50 | 23931336 | 10377.856 |
| HPAP032 | 8705399 | 481 | 26557.00 | 4750158 | 9875.588 |
| HPAP034 | 2320238 | 434 | 6997.25 | 4805112 | 11071.686 |
| HPAP036 | 22928636 | 2597 | 12633.00 | 26300868 | 10127.404 |
| HPAP043 | 4367338 | 1941 | 3102.00 | 20402029 | 10511.092 |
| HPAP048 | 56296579 | 2264 | 34321.75 | 23769001 | 10498.676 |
| HPAP072 | 66625502 | 1052 | 84789.50 | 11386671 | 10823.832 |
| HPAP095 | 32257841 | 1522 | 29219.00 | 15998096 | 10511.233 |
| HPAP102 | 12879491 | 1367 | 12988.00 | 14369934 | 10512.022 |
| HPAP110 | 32190497 | 471 | 121994.00 | 3823733 | 8118.329 |
| HPAP114 | 17730728 | 666 | 44414.50 | 5784957 | 8686.122 |
| HPAP123 | 65698635 | 2125 | 42838.00 | 22224168 | 10458.432 |
| HPAP148 | 6273632 | 3513 | 2018.00 | 4505149 | 12823.840 |
| HPAP149 | 7511295 | 2085 | 4762.00 | 22857240 | 10962.705 |
