## Supplemental Table 2 for "Spatial transcriptomics from pancreas and local draining lymph node tissue reveals a lymphotoxin-β signature in human type 1 diabetes"

**Supplemental Table 3**

Histopathological Notes – pLN

| **CaseID** | **OtherID** | **Review Date** | **Sample** | **#Nodes** | **Complete_All regions** | **Cortex Primary** |
| --- | --- | --- | --- | --- | --- | --- |
| 12 | HPAP | 7/22/24 | PLN | 1 | Yes | Present throughout cortex; not as thick as expected |
| 19 | HPAP | 7/22/24 | PLN | 1 | No | Numerous follicles small-medium |
| 20 | HPAP | 7/22/24 | PLN | 1 | Yes | Numerous follicles medium sized |
| 32 | HPAP | 7/22/24 | PLN | 1 | No | Numerous well-formed med-lg sizes |
| 34 | HPAP | 7/22/24 | PLN | 1 | No | Missing most of cortex, possibly 3 follicles |
| 36 | HPAP | 7/22/24 | PLN | 2 | Yes | Multiple follicles med- lg sized both PLN very well organized |
| 43 | HPAP | 7/22/24 | PLN | 2 | No | Multiple follicles med- lg sized both PLN very well organized |
| 48 | HPAP | 7/22/24 | PLN | 1 | Yes | Follicles not clearly defined. |
| 72 | HPAP | 7/22/24 | PLN | 1 | Yes | Numerous small to medium well organized follicles |
| 95 | HPAP | 7/22/24 | PLN | 1 | Yes | Few medium sized follicles and large flat ones could be plane of sectioning |
| 102 | HPAP | 7/22/24 | PLN | 1 | Yes | Numerous follicles medium to large |
| 110 | HPAP | 7/22/24 | PLN | 1 | No | Numerous follicles small to medium |
| 114 | HPAP | 7/22/24 | PLN | 1 | No | Lacks cortex |
| 123 | HPAP | 7/22/24 | PLN | 1 | Almost (lacks enough medulla) | Numerous small to medium sized well organized |
| 148 | HPAP | 7/22/24 | PLN | 1 | Yes | Numerous med to large sized well organized follicles |
| 149 | HPAP | 7/22/24 | PLN | 1 | Yes | Numerous med to large sized well organized follicles |

| **CaseID** | **Cortex**  **Secondary_GC** | **Medulla** | **Fat_Intra** | **Fibrosis_intra** | **Notes** |
| --- | --- | --- | --- | --- | --- |
| 12 | 2 | Reduced cellularity | 5% | 0 | Good |
| 19 | 0 | WNL | 0% | 0 | Cracking, folds, air bubbles. Lacks hilum. |
| 20 | 1 | Reduced cellularity | 0% | 0 | Good |
| 32 | 0 | WNL small region present | 10% | 0 | Folds, air bubbles; lack hilum and most of medulla |
| 34 | 0 | WNL | 0% | 0 | Folds; missing cortex and hilum |
| 36 | 2 | WNL | 5 (lower node)% | 0 | Few folds both; scrape mark lower node; bubbles outside tissue |
| 43 | 6 | Reduced cellularity | 0% | Grade 2 with sclerosis | Cracking mild, numerous small air bubbles |
| 48 | 0 | WNL | 0% | 0 | Numerous small air bubbles |
| 72 | 3 | WNL | 0% | 0 | Good |
| 95 | 0 | WNL | 0% | 0 | Cracking, knife marks, bubbles |
| 102 | 0 | WNL | 0% | 0 | Cracking, knife marks, bubbles |
| 110 | 0 | WNL | Not enough hilum | Not enough hilum | Folds |
| 114 | NP | Lacks clear cells and morphology secondary to autolysis | NP | Grade 1 | Autolysis moderate to severe with nuclear streaming |
| 123 | 0 | Reduced cellularity but also lacks sufficient area | NP | Grade 2 | Cracking, folds around follicles |
| 148 | 4 | WNL | 0% | 0 | Extremely large node, minor folding and cracks |
| 149 | 9 | WNL | 0% | 0 | Extremely large node, minor folding and cracks |

| **CaseID** | Interlobular fat | **Microadenoma** | Nerve_Prom | **Autolysis (%)** | **Notes** |
| --- | --- | --- | --- | --- | --- |
| 12 | 0 | 0 | 0 | 0 | ND |
| 16 | 2 | 0 | 2 | 0 | ND-like |
| 19 | 1 | 0 | 0 | 0 | ND-like |
| 20 | 0 | 0 | 0 | 0 | T1D |
| 32 | 0 | 0 | 0 | 0 | T1D |
| 34 | 0 | 0 | 0 | 0 | ND |
| 36 | 0 | 0 | 0 | 0 | ND with one foci interstitial fibrosis and very mild exocrine mixed infiltrates |
| 43 | 1 | 0 | 0 | 0 | ND features with no evidence insulitis or exocrine disease |
| 48 | 1 | 0 | 1 | 0 |  |
| 71 | 0 | 0 | 0 | 5 | T1D with small islets |
| 72 | 0 | 0 | 0 | 0 | ND-like |
| 87 | 0 | 0 | 0 | 0 | T1D with few islets |
| 95 | 0 | 0 | 0 | 0 | ND |
| 102 | 0 | 0 | 0 | 0 | T1D with small islets |
| 107 | 1 | 0 | 0 | 0 | ND like but possible insulitic islet |
| 110 | 0 | 0 | 0 | 0 | ND |
| 114 | 1 | 0 | 0 | 0 | ND like with mild fatty infiltration |
| 123 | 0 | 0 | 1 | 0 | T1D like with exocrine infiltrates- islet numbers reduced, acinar atrophy |
| 148 | 0 | 0 | 0 | 0 | ND |
| 149 | 0 | 0 | 0 | 0 | T1D |

**WNL** – Within normal limits

**GC** – Germinal center

**NP** – Not possible
