## Supplemental Table 3 for "Spatial transcriptomics from pancreas and local draining lymph node tissue reveals a lymphotoxin-β signature in human type 1 diabetes"

| **CaseID** | **Islet sizes** | **Islets per section** | **Insulitis** | **Angiopathy (small vessels)** | **Atherosclerosis/Arteriosclerosis** | **Acute inflammation (1 Yes/ 0 No)** |
| --- | --- | --- | --- | --- | --- | --- |
| 12 | Normal range | Numerous | 0 | 0 | 0 | 0 |
| 16 | Normal range |  | 0 | 0 | 1 | 0 |
| 19 | Normal range | Normal | 0 | 0 | 0 | 0 |
| 20 | Reduced | Reduced | 0 | 0 | 0 | 0 |
| 32 | Reduced | Reduced | 0 | 0 | 0 | 0.5- possible mild global mixed WBC MN infil |
| 34 | Normal range | Numerous | 0 | 0 | 0 | 0 |
| 36 | Normal range | Numerous | 0 | 0 | 0 | 0.5- possible mild global mixed WBC MN infil |
| 43 | Normal range | Numerous | 0 | 0 | 0 | 0 |
| 48 | Medium | Reduced | 0 | 0 | 2 | 0 |
| 71 | Small | Reduced | 0 | 0 | 0 | 0 |
| 72 | Normal range | Normal | 0 | 0 | 0 | 0 |
| 87 | Small | Reduced | 0 | 0 | 0 | 0 |
| 95 | Normal range | Numerous | 0 | 0 | 0 | 0 |
| 102 | Small | Reduced | 0 | 0 | 0 | 0 |
| 107 | Normal range | Numerous | 1 islet possible [boxed] | 0 | 0 | 0 |
| 110 | Normal range | Numerous | 0 | 0 | 1 | 0 |
| 114 | Normal range | Numerous | 0 | 0 | 0 | 0 |
| 123 | Small mostly | Reduced | 0 | 0 | 0 | 0 |
| 148 | Normal range | Numerous | 0 | 0 | 0 | 0 |
| 149 | Too few to describe | greatly reduced | not possible | 0 | 0 | 0 |

**Supplemental Table 3**

Histopathological Notes – Pancreas

| **CaseID** | Chronic inflammation (1 Yes/ 0 No) | Periductal MN infiltration | Acinar atrophy | Intralobular  Fibrosis | Interlobular Fibrosis | Interstitial  Fibrosis | **Duct**  **Proliferation** | **Duct**  **Dilation** | Intralobular fat |
| --- | --- | --- | --- | --- | --- | --- | --- | --- | --- |
| 12 | 0 | 0 | 0 | 0 | 0 | 0 | 0 | 0 | 0 |
| 16 | 2 | 2 | 0 | 2 | 2 | 2 | 2 | 0 | 2 |
| 19 | 0 | 0 | 0 | 0 | 0 | 0 | 0 | 0 | 0 |
| 20 | 0.5 | 0 | 1 | 0 | 1 | 0 | 0 | 0 | 0 |
| 32 | 0.5- mild global mixed WBC MN infil | 0 | 1 | 0 | 1 | 0 | 0 | 0 | 0 |
| 34 | 0 | 0 | 0 | 0 | 0 | 0 | 0 | 0 | 0 |
| 36 | 0 | 0 | 1 | 0 | 1 | 1 | 0 | 0 | 0 |
| 43 | 0 | 0 | 0 | 0 | 0 | 0 | 0 | 0 | 0 |
| 48 | 0 | 0 | 1 | 1 | 1 | 0 | 0 | 0 | 0 |
| 71 | 0 | 0 | 0.5 (periductal regions) | 1 | 0 | 0 | 0 | 0 | 0 |
| 72 | 0 | 0 | 0 | 0 | 0 | 0 | 0 | 0 | 0 |
| 87 | 0 | 0 | 1 (periductal regions) | 0 | 0 | 0 | 0 | 0 | 0 |
| 95 | 0 | 0 | 0 | 0 | 0 | 0 | 0 | 0 | 0 |
| 102 | 0 | 0 | 1 | 1 | 1 | 0 | 0 | 0 | 0 |
| 107 | 0.5- periphery lobule mononuclear | 0 | 0 | 0 | 0 | 0 | 0 | 0 | 0 |
| 110 | 0 | 0 | 1 | 0.5 | 0 | 1 | 0 | Very early ADM | 1 |
| 114 | 0 | 0 | 1 | 0 | 1 | 0 | 0 | 0 | 2 |
| 123 | 1 (multi focal primarily MN infiltration interlobular extending along fascial planes) |  | 1 | 1 | 0 | 0 | 0 | 0 | 0 |
| 148 | 0 | 0 | 0 | 0 | 0 | 0 | 0 | 0 | 0 |
| 149 | 0 | 0 | 1 | 0.5 | 0 | 0 | 0 | 0 | 0 |

| **CaseID** | Interlobular fat | **Microadenoma** | Nerve_Prom | **Autolysis %** | **Notes** |
| --- | --- | --- | --- | --- | --- |
| 12 | 0 | 0 | 0 | 0 | ND |
| 16 | 2 | 0 | 2 | 0 | ND-like |
| 19 | 1 | 0 | 0 | 0 | ND-like |
| 20 | 0 | 0 | 0 | 0 | T1D |
| 32 | 0 | 0 | 0 | 0 | T1D |
| 34 | 0 | 0 | 0 | 0 | ND |
| 36 | 0 | 0 | 0 | 0 | ND with one foci interstital fibrosis and very mild exocrine mixed infiltrates |
| 43 | 1 | 0 | 0 | 0 | ND features with no evidence insulitis or exocrine disease |
| 48 | 1 | 0 | 1 | 0 |  |
| 71 | 0 | 0 | 0 | 5 | T1D with small islets |
| 72 | 0 | 0 | 0 | 0 | ND-like |
| 87 | 0 | 0 | 0 | 0 | T1D with few islets |
| 95 | 0 | 0 | 0 | 0 | ND |
| 102 | 0 | 0 | 0 | 0 | T1D with small islets |
| 107 | 1 | 0 | 0 | 0 | ND like but possible insulitic islet |
| 110 | 0 | 0 | 0 | 0 | ND |
| 114 | 1 | 0 | 0 | 0 | ND like with mild fatty infiltration |
| 123 | 0 | 0 | 1 | 0 | T1D like with exocrine infiltrates- islet numbers reduced, acinar atrophy |
| 148 | 0 | 0 | 0 | 0 | ND |
| 149 | 0 | 0 | 0 | 0 | T1D |

**WBC –** White blood cells

**MN** – Mononuclear

**ADM** – Acinar-to-ductal metaplasia
