## Supplemental Table 4 for "Spatial transcriptomics from pancreas and local draining lymph node tissue reveals a lymphotoxin-β signature in human type 1 diabetes"

**Supplemental Table 8**

Regional differential expression – Endocrine (Pancreas)

|  | Gene | p_val | avg_log2FC | rank | BH.FDR |
| --- | --- | --- | --- | --- | --- |
| **Endocrine - AAb+ vs ND** | REG1B | 8.16E-23 | -0.6217125 | 1 | **8.16E-20** |
|  | REG3A | 3.27E-20 | **-1.6475475** | 2 | **1.63E-17** |
|  | REG3G | 1.56E-13 | **-1.6670049** | 3 | **5.20E-11** |
|  | INS | 5.26E-11 | -0.3977479 | 4 | **1.32E-08** |
|  | CTRB2 | 2.53E-08 | -0.3702062 | 5 | **5.06E-06** |
|  | CELA2B | 3.56E-07 | 0.85413684 | 6 | **5.93E-05** |
|  | GCG | 1.21E-06 | **-1.1187407** | 7 | **0.00017291** |
|  | CUZD1 | 1.53E-06 | **-1.0031864** | 8 | **0.00019097** |
|  | TTR | 2.95E-06 | **-1.1213249** | 9 | **0.00032759** |
|  | SERPINA3 | 9.98E-05 | **-1.1800219** | 10 | **0.00998038** |
|  | OLFM4 | 0.00034264 | **-2.131046** | 11 | **0.03114904** |
|  | PPY | 0.00124404 | **-4.1490806** | 12 | 0.10367024 |
|  | SST | 0.00240735 | -0.7698316 | 13 | 0.18518067 |
|  | IAPP | 0.00321332 | -1.0610229 | 14 | 0.22952286 |
|  | SOD2 | 0.01578156 | -1.1623333 | 15 | 1 |
|  | RNASE1 | 0.02101562 | 0.4734784 | 16 | 1 |
|  | CHGB | 0.02383422 | -1.0239477 | 17 | 1 |
|  | AMY2A | 0.02509852 | 0.53712084 | 18 | 1 |
|  | CEBPD | 0.02595428 | -1.1181865 | 19 | 1 |

|  | Gene | p_val | avg_log2FC | rank | BH.FDR |
| --- | --- | --- | --- | --- | --- |
| **Endocrine - T1D vs ND** | INS | 2.13E-129 | **-6.347994** | 1 | **2.13E-126** |
|  | REG1B | 5.32E-105 | 0.84661461 | 2 | **2.66E-102** |
|  | REG3A | 1.09E-55 | **1.26914519** | 3 | **3.62E-53** |
|  | OLFM4 | 7.05E-53 | **3.32661499** | 4 | **1.76E-50** |
|  | CTRB2 | 5.96E-41 | -0.7132338 | 5 | **1.19E-38** |
|  | C3 | 1.95E-24 | **2.32648386** | 6 | **3.24E-22** |
|  | CELA2A | 3.51E-13 | -0.7330389 | 7 | **5.02E-11** |
|  | PPY | 7.60E-12 | **1.09886967** | 8 | **9.50E-10** |
|  | ALDOB | 3.73E-09 | **1.84301847** | 9 | **4.15E-07** |
|  | SOD2 | 2.58E-07 | **1.33828691** | 10 | **2.58E-05** |
|  | SERPINA3 | 8.15E-07 | 0.80628355 | 11 | **7.41E-05** |
|  | MT1X | 9.34E-07 | **1.39514692** | 12 | **7.78E-05** |
|  | IAPP | 1.05E-06 | **-3.5006652** | 13 | **8.07E-05** |
|  | SERPINB1 | 5.70E-06 | **1.10543223** | 14 | **0.00040706** |
|  | RNASE1 | 8.45E-05 | -0.773798 | 15 | **0.00563429** |
|  | CD44 | 9.48E-05 | **1.1258563** | 16 | **0.00592642** |
|  | IL32 | 0.00020099 | 0.99351003 | 17 | **0.01182312** |
|  | PDK4 | 0.00022617 | **1.14231765** | 18 | **0.01256518** |
|  | CELA2B | 0.00026459 | -0.6113208 | 19 | **0.01392576** |
|  | TMSB4X | 0.00066478 | 0.6009004 | 20 | **0.03165601** |
|  | TMSB10 | 0.00079308 | 0.94981543 | 21 | **0.03604885** |
|  | G6PC2 | 0.00091906 | -1.0673546 | 22 | **0.03995903** |
|  | TPM1 | 0.00102931 | 0.97524835 | 23 | **0.04288791** |
|  | B2M | 0.00118012 | 0.7504644 | 24 | **0.04720473** |
|  | REG3G | 0.00231179 | 0.33563081 | 25 | 0.08891504 |
|  | LYZ | 0.00253062 | 1.38459333 | 26 | 0.09372678 |
|  | MMP7 | 0.00301956 | 1.29676669 | 27 | 0.10784128 |
|  | CPE | 0.0042774 | -0.800612 | 28 | 0.14749645 |
|  | ACTB | 0.00492204 | 0.89385217 | 29 | 0.16406784 |
|  | TACSTD2 | 0.00624461 | 1.01015217 | 30 | 0.20143891 |
|  | DEPP1 | 0.00703099 | 0.82292839 | 31 | 0.21971853 |
|  | HSPB1 | 0.00774387 | 0.75742224 | 32 | 0.23466264 |
|  | TIMP1 | 0.01124597 | 1.02383326 | 33 | 0.33076387 |
|  | CUZD1 | 0.01217826 | -0.3357092 | 34 | 0.34795039 |
|  | PPP1R1A | 0.01479504 | -1.1104407 | 35 | 0.41097335 |
|  | ADAMTS1 | 0.02671309 | 1.05624903 | 36 | 0.72197534 |
|  | C11orf96 | 0.03474953 | 0.99688604 | 37 | 0.91446121 |
|  | KRT7 | 0.04433191 | 0.86689757 | 38 | 1 |
|  | SPP1 | 0.04569134 | 0.71336291 | 39 | 1 |
|  | PTP4A3 | 0.04775421 | 0.73303136 | 40 | 1 |

| **Endocrine - T1D vs AAb+** | Gene | p_val | avg_log2FC | rank | BH.FDR |
| --- | --- | --- | --- | --- | --- |
|  | REG1B | 2.48E-145 | **1.46832707** | 1 | **2.48E-142** |
|  | INS | 1.04E-112 | **-5.9502461** | 2 | **5.21E-110** |
|  | REG3A | 1.52E-66 | **2.91669264** | 3 | **5.05E-64** |
|  | REG3G | 3.50E-20 | **2.00263576** | 4 | **8.76E-18** |
|  | CELA2B | 7.78E-19 | **-1.4654576** | 5 | **1.56E-16** |
|  | C3 | 1.74E-16 | **2.79248577** | 6 | **2.59E-14** |
|  | OLFM4 | 1.81E-16 | **5.45766099** | 7 | **2.59E-14** |
|  | SERPINA3 | 5.04E-12 | **1.9863055** | 8 | **6.30E-10** |
|  | CELA2A | 2.91E-10 | -0.7316709 | 9 | **3.23E-08** |
|  | RNASE1 | 1.71E-09 | **-1.2472764** | 10 | **1.71E-07** |
|  | TTR | 2.69E-08 | **1.27363208** | 11 | **2.45E-06** |
|  | AMY2A | 4.12E-08 | **-1.3294631** | 12 | **3.44E-06** |
|  | SOD2 | 2.21E-07 | **2.5006202** | 13 | **1.70E-05** |
|  | CTRB2 | 2.66E-07 | -0.3430276 | 14 | **1.90E-05** |
|  | SERPINB1 | 1.40E-06 | **1.83011097** | 15 | **9.31E-05** |
|  | ALDOB | 1.74E-06 | **1.95127465** | 16 | **0.0001089** |
|  | GCG | 2.94E-06 | **1.03735344** | 17 | **0.00017281** |
|  | MT1X | 5.92E-06 | **1.89436111** | 18 | **0.00032908** |
|  | CD44 | 9.00E-05 | **1.66128923** | 19 | **0.00473706** |
|  | IAPP | 0.00012798 | **-2.4396422** | 20 | **0.00639885** |
|  | PPY | 0.00017254 | **5.24795031** | 21 | **0.00821633** |
|  | PDK4 | 0.00023375 | **1.6924888** | 22 | **0.01062476** |
|  | TMSB10 | 0.0003133 | **1.48513951** | 23 | **0.01362159** |
|  | CUZD1 | 0.00100436 | 0.66747716 | 24 | **0.04184832** |
|  | ACTB | 0.00107509 | **1.62050258** | 25 | **0.0430035** |
|  | IL32 | 0.00153553 | 1.06534601 | 26 | 0.05905897 |
|  | DEPP1 | 0.00184784 | 1.39334928 | 27 | 0.06843843 |
|  | CHGB | 0.00199413 | 1.35466529 | 28 | 0.07121901 |
|  | MUC6 | 0.00342642 | 1.06191527 | 29 | 0.11815239 |
|  | CEBPD | 0.004457 | 1.39484136 | 30 | 0.14856661 |
|  | HSPB1 | 0.0066269 | 1.02095695 | 31 | 0.2137708 |
|  | DEFB1 | 0.00695679 | 1.62440324 | 32 | 0.21739982 |
|  | LYZ | 0.01072305 | 1.64958358 | 33 | 0.32494077 |
|  | JUNB | 0.01175676 | 1.50164282 | 34 | 0.34578713 |
|  | B2M | 0.01217989 | 1.13425428 | 35 | 0.34799682 |
|  | MMP7 | 0.01258689 | 1.44829427 | 36 | 0.34963572 |
|  | TIMP1 | 0.01385622 | 1.42246547 | 37 | 0.3744923 |
|  | TACSTD2 | 0.01498963 | 1.1708204 | 38 | 0.39038941 |
|  | PTP4A3 | 0.01535383 | 1.34792003 | 39 | 0.39038941 |
|  | SST | 0.01561558 | 0.58845082 | 40 | 0.39038941 |
|  | MT2A | 0.02136269 | 1.12007993 | 41 | 0.52104127 |
|  | S100A6 | 0.02387797 | 1.30813571 | 42 | 0.56269122 |
|  | TPM1 | 0.02419572 | 0.77880596 | 43 | 0.56269122 |
|  | SPP1 | 0.02836855 | 1.06860233 | 44 | 0.63460982 |
|  | TXNIP | 0.02855744 | 1.05181955 | 45 | 0.63460982 |
|  | IFITM3 | 0.03398045 | 1.27727103 | 46 | 0.73870549 |
|  | KRT7 | 0.0396213 | 1.29214315 | 47 | 0.84300641 |
