## Supplemental Table 5 for "Spatial transcriptomics from pancreas and local draining lymph node tissue reveals a lymphotoxin-β signature in human type 1 diabetes"

**Supplemental Table 9**

Regional differential expression – Exocrine (Pancreas)

|  | Gene | p_val | avg_log2FC | rank | BH.FDR |
| --- | --- | --- | --- | --- | --- |
| **Exocrine**  **AAb+ vs ND** | REG3A | 8.49E-30 | **-1.6972851** | 1 | 8.49E-27 |
|  | REG3G | 5.04E-17 | **-1.5097367** | 2 | 2.52E-14 |
|  | CTRB2 | 7.18E-11 | -0.3774332 | 3 | 2.39E-08 |
|  | OLFM4 | 5.47E-06 | **-1.7468428** | 4 | 0.00136841 |
|  | PPY | 0.02390512 | -1.6700651 | 5 | 1 |
|  | CELA2B | 0.02510033 | 0.34883532 | 6 | 1 |
|  | CELA2A | 0.02688681 | -0.215198 | 7 | 1 |
|  | CEBPD | 0.02870396 | -0.8232365 | 8 | 1 |
|  | GNMT | 0.04223144 | -0.6014222 | 9 | 1 |
|  | CLPSL1 | 0.04833895 | 0.88631412 | 10 | 1 |

|  | Gene | p_val | avg_log2FC | rank | BH.FDR |
| --- | --- | --- | --- | --- | --- |
| **Exocrine - T1D vs ND** | REG1B | 1.16E-142 | **1.03466529** | 1 | 1.16E-139 |
|  | REG3A | 7.34E-90 | **1.71725129** | 2 | 3.67E-87 |
|  | OLFM4 | 3.13E-61 | **3.58191364** | 3 | 1.04E-58 |
|  | C3 | 1.14E-27 | **2.63108246** | 4 | 2.84E-25 |
|  | CTRB2 | 8.76E-16 | -0.4386775 | 5 | 1.75E-13 |
|  | CELA2A | 1.54E-14 | -0.7639153 | 6 | 2.56E-12 |
|  | SERPINA3 | 3.82E-11 | **1.11249599** | 7 | 5.46E-09 |
|  | SOD2 | 9.31E-10 | **1.63050596** | 8 | 1.03E-07 |
|  | REG3G | 1.88E-08 | 0.6555125 | 9 | 1.88E-06 |
|  | ALDOB | 2.58E-07 | **1.64588496** | 10 | 2.35E-05 |
|  | MT1X | 1.92E-06 | **1.3381486** | 11 | 0.00015967 |
|  | SERPINB1 | 2.21E-06 | **1.16300943** | 12 | 0.00016963 |
|  | IL32 | 1.02E-05 | **1.21963298** | 13 | 0.00072846 |
|  | CD44 | 1.39E-05 | **1.31754984** | 14 | 0.00092903 |
|  | TMSB4X | 0.00012292 | 0.72232907 | 15 | 0.00768262 |
|  | TPM1 | 0.0001494 | **1.18166352** | 16 | 0.00878843 |
|  | PDK4 | 0.00046737 | **1.13501145** | 17 | 0.0259649 |
|  | CELA2B | 0.00053083 | -0.5794478 | 18 | 0.02793825 |
|  | RNASE1 | 0.00066694 | -0.6829716 | 19 | 0.03334713 |
|  | TMSB10 | 0.00075774 | 0.99819857 | 20 | 0.03608307 |
|  | HSPB1 | 0.00081162 | 0.96076979 | 21 | 0.03689192 |
|  | MMP7 | 0.00097609 | **1.45378102** | 22 | 0.04243851 |
|  | DEPP1 | 0.00143051 | 1.06769647 | 23 | 0.05960459 |
|  | AMY2A | 0.00197044 | -0.687673 | 24 | 0.07881772 |
|  | TACSTD2 | 0.00362118 | 1.09341635 | 25 | 0.13927622 |
|  | ACTB | 0.004477 | 0.95039224 | 26 | 0.16581468 |
|  | SPP1 | 0.00481456 | 1.10470304 | 27 | 0.17194853 |
|  | LYZ | 0.00506797 | 1.30584484 | 28 | 0.17475744 |
|  | B2M | 0.00860061 | 0.94399292 | 29 | 0.28668694 |
|  | TIMP1 | 0.01293029 | 1.12568666 | 30 | 0.41710624 |
|  | PPY | 0.02323549 | 0.83315383 | 31 | 0.72610919 |
|  | SAT1 | 0.02582094 | 0.92349165 | 32 | 0.7671549 |
|  | PTP4A3 | 0.02608327 | 0.87728883 | 33 | 0.7671549 |
|  | IFI6 | 0.02748142 | 1.17075343 | 34 | 0.78518347 |
|  | MT2A | 0.0287955 | 0.78324805 | 35 | 0.79987491 |
|  | ADAMTS1 | 0.03114594 | 1.02619195 | 36 | 0.84178209 |
|  | C11orf96 | 0.03735097 | 0.99389027 | 37 | 0.98292037 |
|  | DUSP23 | 0.04074134 | 0.987329 | 38 | 1 |
|  | S100A6 | 0.04597621 | 0.82723436 | 39 | 1 |

| **Exocrine - T1D vs AAb+** | Gene | p_val | avg_log2FC | rank | BH.FDR |
| --- | --- | --- | --- | --- | --- |
|  | REG1B | 2.06E-172 | **1.07446851** | 1 | 2.06E-169 |
|  | REG3A | 4.31E-143 | **3.4145364** | 2 | 2.16E-140 |
|  | OLFM4 | 3.28E-42 | **5.32875648** | 3 | 1.09E-39 |
|  | REG3G | 1.95E-39 | **2.16524924** | 4 | 4.87E-37 |
|  | C3 | 1.99E-32 | **3.19615826** | 5 | 3.98E-30 |
|  | SERPINA3 | 6.77E-16 | **1.34935564** | 6 | 1.13E-13 |
|  | SOD2 | 2.10E-13 | **2.08227552** | 7 | 3.00E-11 |
|  | CELA2B | 1.07E-09 | -0.9282831 | 8 | 1.33E-07 |
|  | MT1X | 3.90E-09 | **1.71668748** | 9 | 3.90E-07 |
|  | CELA2A | 2.59E-08 | -0.5487173 | 10 | 2.35E-06 |
|  | AMY2A | 2.55E-07 | **-1.049426** | 11 | 2.12E-05 |
|  | ALDOB | 1.21E-06 | **1.29920389** | 12 | 9.30E-05 |
|  | SERPINB1 | 1.47E-06 | **1.07752148** | 13 | 0.00010533 |
|  | IL32 | 3.11E-06 | **1.18974705** | 14 | 0.00020734 |
|  | TMSB10 | 3.85E-06 | **1.44187962** | 15 | 0.0002406 |
|  | RNASE1 | 9.05E-06 | -0.8361198 | 16 | 0.0005322 |
|  | PDK4 | 1.69E-05 | **1.39860047** | 17 | 0.00093819 |
|  | HSPB1 | 3.58E-05 | **1.16797326** | 18 | 0.00188578 |
|  | DEPP1 | 0.00015377 | **1.23547374** | 19 | 0.00768823 |
|  | TMSB4X | 0.0003899 | 0.60888291 | 20 | 0.01856676 |
|  | CEBPD | 0.00047945 | **1.21611038** | 21 | 0.02124241 |
|  | ACTB | 0.00048858 | **1.14440066** | 22 | 0.02124241 |
|  | TACSTD2 | 0.00055037 | **1.26955709** | 23 | 0.02293218 |
|  | MMP7 | 0.0006202 | **1.41618558** | 24 | 0.02480807 |
|  | B2M | 0.00089124 | **1.18878575** | 25 | 0.03427833 |
|  | TPM1 | 0.00147457 | 0.84571389 | 26 | 0.05461387 |
|  | LYZ | 0.0015541 | 1.42385172 | 27 | 0.05550361 |
|  | PTP4A3 | 0.00169829 | 1.3044275 | 28 | 0.05856176 |
|  | DEFB1 | 0.00178419 | 1.20728591 | 29 | 0.05947289 |
|  | TIMP1 | 0.00232848 | 1.42449511 | 30 | 0.07511222 |
|  | CD44 | 0.0024523 | 0.73483762 | 31 | 0.07663421 |
|  | MT2A | 0.00278543 | 1.08015968 | 32 | 0.08440706 |
|  | JUNB | 0.00320956 | 1.18946116 | 33 | 0.09439868 |
|  | GNMT | 0.00362335 | 0.77975111 | 34 | 0.10352427 |
|  | MUC6 | 0.00538054 | 0.70529738 | 35 | 0.14795419 |
|  | SPP1 | 0.00554939 | 0.97571958 | 36 | 0.14795419 |
|  | PPY | 0.00562226 | 2.50321897 | 37 | 0.14795419 |
|  | S100A6 | 0.00605636 | 1.16893298 | 38 | 0.15529124 |
|  | CLPSL1 | 0.00726237 | -1.057566 | 39 | 0.18155927 |
|  | ADAMTS1 | 0.00977848 | 1.23918351 | 40 | 0.23845072 |
|  | SERPING1 | 0.01001493 | 1.01010839 | 41 | 0.23845072 |
|  | SLC30A2 | 0.01218369 | 1.07246394 | 42 | 0.28181014 |
|  | C11orf96 | 0.01252881 | 1.19627916 | 43 | 0.28181014 |
|  | TXNIP | 0.01268146 | 0.87693303 | 44 | 0.28181014 |
|  | IFITM3 | 0.01326935 | 1.00673385 | 45 | 0.28846418 |
|  | SAT1 | 0.01544388 | 0.93758566 | 46 | 0.32859319 |
|  | IFI6 | 0.01963762 | 1.10517869 | 47 | 0.4091171 |
|  | KRT7 | 0.02061394 | 1.00906806 | 48 | 0.41373454 |
|  | DUSP23 | 0.02068673 | 1.06338456 | 49 | 0.41373454 |
|  | TM4SF1 | 0.02707044 | 0.93917805 | 50 | 0.53079295 |
|  | SERPINA1 | 0.03028246 | 0.96053346 | 51 | 0.5823549 |
|  | CCL2 | 0.0340963 | 0.93748756 | 52 | 0.6433265 |
|  | FGL1 | 0.03574991 | 0.68699853 | 53 | 0.66203545 |
|  | AHNAK | 0.03786268 | 0.84797257 | 54 | 0.68841241 |
|  | MYL6 | 0.03860344 | 0.72513912 | 55 | 0.68934721 |
|  | RHOB | 0.03968839 | 0.83172463 | 56 | 0.69628754 |
|  | TTR | 0.04431024 | 0.77340746 | 57 | 0.75704842 |
|  | CXCL2 | 0.04466586 | 0.83158583 | 58 | 0.75704842 |
|  | VWA1 | 0.04852228 | 0.86390094 | 59 | 0.80870463 |
