## Supplemental Table 6 for "Spatial transcriptomics from pancreas and local draining lymph node tissue reveals a lymphotoxin-β signature in human type 1 diabetes"

**Supplemental Table 10**

Regional differential expression – Stroma (Pancreas)

|  | Gene | p_val | avg_log2FC | rank | BH.FDR |
| --- | --- | --- | --- | --- | --- |
| **Stroma - AAb+ vs ND** | REG3A | 2.14E-16 | **-1.2681832** | 1 | 2.14E-13 |
|  | REG3G | 2.04E-05 | -0.8747544 | 2 | 0.01019903 |
|  | REG1B | 6.92E-05 | 0.24223677 | 3 | 0.01730503 |
|  | CELA2B | 6.92E-05 | 0.7092213 | 4 | 0.01730503 |
|  | AMY2A | 0.00198598 | 0.69597611 | 5 | 0.39719528 |
|  | OLFM4 | 0.00337246 | -1.2668421 | 6 | 0.56207585 |
|  | PPY | 0.0078121 | -1.8684619 | 7 | 1 |
|  | JUNB | 0.01293093 | -1.0310258 | 8 | 1 |
|  | CEBPD | 0.03201826 | -0.8029395 | 9 | 1 |
|  | INS | 0.03227865 | -0.4250563 | 10 | 1 |
|  | MYL9 | 0.0343751 | -0.9323484 | 11 | 1 |
|  | C11orf96 | 0.04606714 | -0.8749561 | 12 | 1 |
|  | ZFP36 | 0.04758136 | -0.8752273 | 13 | 1 |
|  | RHOB | 0.04859568 | -0.8040517 | 14 | 1 |

|  | Gene | p_val | avg_log2FC | rank | BH.FDR |
| --- | --- | --- | --- | --- | --- |
| **Stroma - T1D vs ND** | REG1B | 1.00E-50 | 0.81375987 | 1 | 1.00E-47 |
|  | REG3A | 1.34E-37 | **1.28910275** | 2 | 6.68E-35 |
|  | OLFM4 | 5.78E-30 | **3.40114129** | 3 | 1.93E-27 |
|  | CTRB2 | 6.21E-18 | -0.5823723 | 4 | 1.55E-15 |
|  | CELA2A | 1.67E-14 | -1.009696 | 5 | 3.35E-12 |
|  | C3 | 3.77E-13 | **1.78642712** | 6 | 5.39E-11 |
|  | SOD2 | 3.52E-06 | **1.38925383** | 7 | 0.00044036 |
|  | CELA2B | 5.37E-05 | -0.8390617 | 8 | 0.00596384 |
|  | SERPINA3 | 0.00099777 | 0.68415214 | 9 | 0.09422282 |
|  | IL32 | 0.00103645 | 1.08831365 | 10 | 0.09422282 |
|  | CUZD1 | 0.00129042 | -0.5781817 | 11 | 0.10753529 |
|  | CD44 | 0.00266453 | 1.05015297 | 12 | 0.19672109 |
|  | TMSB4X | 0.0027541 | 0.59515844 | 13 | 0.19672109 |
|  | TMSB10 | 0.00295667 | 0.90415813 | 14 | 0.19711141 |
|  | SERPINB1 | 0.0037553 | 0.89545946 | 15 | 0.23470617 |
|  | MT1X | 0.00469776 | 0.8995933 | 16 | 0.27633851 |
|  | RNASE1 | 0.00610782 | -0.6634136 | 17 | 0.33734976 |
|  | AMY2A | 0.00640965 | -0.6888599 | 18 | 0.33734976 |
|  | MMP7 | 0.0106447 | 1.19317349 | 19 | 0.53223504 |
|  | TIMP1 | 0.01334926 | 0.97721552 | 20 | 0.61737711 |
|  | SPP1 | 0.0135823 | 1.02230473 | 21 | 0.61737711 |
|  | HSPB1 | 0.0165729 | 0.71955969 | 22 | 0.72056068 |
|  | ALDOB | 0.01880246 | 0.84944211 | 23 | 0.76490619 |
|  | ACTB | 0.01912266 | 0.74916425 | 24 | 0.76490619 |
|  | REG3G | 0.02694515 | 0.35176399 | 25 | 1 |
|  | TPM1 | 0.032221 | 0.69927876 | 26 | 1 |
|  | VIM | 0.03586258 | 0.47068664 | 27 | 1 |
|  | DEPP1 | 0.04728075 | 0.74356209 | 28 | 1 |

|  | Gene | p_val | avg_log2FC | rank | BH.FDR |
| --- | --- | --- | --- | --- | --- |
| **Stroma - T1D vs AAb+** | REG3A | 2.12E-79 | **2.55728595** | 1 | 2.12E-76 |
|  | REG1B | 1.23E-32 | 0.5715231 | 2 | 6.17E-30 |
|  | OLFM4 | 1.69E-23 | **4.66798338** | 3 | 5.64E-21 |
|  | CTRB2 | 1.02E-21 | -0.6146226 | 4 | 2.56E-19 |
|  | C3 | 1.32E-17 | **2.23555315** | 5 | 2.63E-15 |
|  | CELA2B | 5.40E-17 | **-1.548283** | 6 | 9.00E-15 |
|  | CELA2A | 7.90E-17 | **-1.0501673** | 7 | 1.13E-14 |
|  | REG3G | 3.27E-11 | **1.22651841** | 8 | 3.64E-09 |
|  | AMY2A | 7.39E-10 | **-1.384836** | 9 | 7.39E-08 |
|  | SOD2 | 7.71E-09 | **1.8385646** | 10 | 7.01E-07 |
|  | RNASE1 | 1.45E-06 | **-1.064686** | 11 | 0.00012043 |
|  | SERPINA3 | 6.40E-06 | 0.93423484 | 12 | 0.00049233 |
|  | TMSB10 | 5.42E-05 | **1.25905086** | 13 | 0.0038739 |
|  | ACTB | 0.00038835 | **1.1872553** | 14 | 0.02589003 |
|  | TIMP1 | 0.00048332 | **1.51933773** | 15 | 0.03020738 |
|  | MT1X | 0.00074434 | **1.03793139** | 16 | 0.04378494 |
|  | CUZD1 | 0.00121354 | -0.5577243 | 17 | 0.06407006 |
|  | TAGLN | 0.001233 | 1.17548637 | 18 | 0.06407006 |
|  | S100A6 | 0.0012814 | 1.21730331 | 19 | 0.06407006 |
|  | HSPB1 | 0.00162046 | 0.93345406 | 20 | 0.07716461 |
|  | SERPINB1 | 0.00300215 | 0.84586619 | 21 | 0.13646125 |
|  | C11orf96 | 0.00412275 | 1.20953701 | 22 | 0.17924983 |
|  | VIM | 0.00477031 | 0.61467388 | 23 | 0.19876284 |
|  | MYL9 | 0.00521894 | 1.18058205 | 24 | 0.20546289 |
|  | PPY | 0.00534204 | 2.00398524 | 25 | 0.20546289 |
|  | B2M | 0.00604049 | 0.98278456 | 26 | 0.22372191 |
|  | PDK4 | 0.0075016 | 0.88915493 | 27 | 0.26791434 |
|  | IL32 | 0.00835924 | 0.74817155 | 28 | 0.2865497 |
|  | LYZ | 0.00859649 | 1.09809209 | 29 | 0.2865497 |
|  | MUC6 | 0.00911915 | 0.70760372 | 30 | 0.29416596 |
|  | ADAMTS1 | 0.00954489 | 1.13134925 | 31 | 0.2982777 |
|  | JUNB | 0.01086143 | 0.99408966 | 32 | 0.3291342 |
|  | TPM2 | 0.0144399 | 1.04048064 | 33 | 0.414144 |
|  | MMP7 | 0.01449504 | 0.99413118 | 34 | 0.414144 |
|  | THBS1 | 0.0182398 | 1.0084042 | 35 | 0.50666096 |
|  | MYH11 | 0.01918687 | 1.11722548 | 36 | 0.51856403 |
|  | CEBPD | 0.02113156 | 0.80181861 | 37 | 0.55609373 |
|  | TMSB4X | 0.02376545 | 0.40438793 | 38 | 0.60937049 |
|  | CLPSL1 | 0.0252199 | -0.9317454 | 39 | 0.63049751 |
|  | DES | 0.02964695 | 1.46123914 | 40 | 0.72309644 |
|  | CD44 | 0.03309768 | 0.62538467 | 41 | 0.78804003 |
|  | MT2A | 0.03677283 | 0.74976538 | 42 | 0.85518212 |
|  | IGFBP4 | 0.03921128 | 0.7853095 | 43 | 0.89116546 |
|  | DEPP1 | 0.04074468 | 0.71136022 | 44 | 0.8979584 |
|  | IFITM3 | 0.04172362 | 0.81832622 | 45 | 0.8979584 |
|  | TACSTD2 | 0.04220405 | 0.77291342 | 46 | 0.8979584 |
|  | COL6A3 | 0.04334086 | 0.77946507 | 47 | 0.90293464 |
|  | AHNAK | 0.04685537 | 0.74206316 | 48 | 0.95623193 |
|  | ACTA2 | 0.04803531 | 0.88699309 | 49 | 0.96070624 |
