## Supplemental Table 7 for "Spatial transcriptomics from pancreas and local draining lymph node tissue reveals a lymphotoxin-β signature in human type 1 diabetes"

|  | gene | p_val | avg_log2FC | rank | BH.FDR |
| --- | --- | --- | --- | --- | --- |
| **Lymphoid Follicle - AAb+ vs ND** | IGKC | 8.47E-80 | **-1.6009314** | 1 | 4.23E-77 |
|  | IGHM | 5.39E-14 | -0.9922327 | 2 | 1.35E-11 |
|  | IGHA1 | 2.55E-13 | **-1.7271436** | 3 | 4.26E-11 |
|  | IGHG1 | 6.96E-08 | -0.9400111 | 4 | 8.70E-06 |
|  | JCHAIN | 4.80E-06 | **-1.3032345** | 5 | 0.00047963 |
|  | CD52 | 6.52E-06 | **-1.1576005** | 6 | 0.00054324 |
|  | TXNDC5 | 0.00027631 | **-1.4512045** | 7 | 0.01973613 |
|  | P2RX5 | 0.00027766 | **-1.0124015** | 8 | 0.01735393 |
|  | CD79A | 0.00184943 | -0.7287052 | 9 | 0.10274582 |
|  | FCMR | 0.00429719 | -0.8227636 | 10 | 0.21485959 |
|  | IGLC1 | 0.00529449 | -1.1731917 | 11 | 0.24065866 |
|  | CCN1 | 0.00537866 | -1.1687027 | 12 | 0.22411093 |
|  | BANK1 | 0.00709873 | -0.7703181 | 13 | 0.27302812 |
|  | TNFRSF13C | 0.00766621 | -0.7149988 | 14 | 0.27379326 |
|  | CD37 | 0.00964794 | -0.5220105 | 15 | 0.32159802 |
|  | STK17B | 0.01740745 | -0.7479002 | 16 | 0.54398293 |
|  | TMC8 | 0.02233682 | -0.6460173 | 17 | 0.65696518 |
|  | EGR1 | 0.02267939 | -0.7768524 | 18 | 0.62998299 |
|  | CD79B | 0.02308521 | -0.7362632 | 19 | 0.60750562 |
|  | CLU | 0.02943831 | 0.63492161 | 20 | 0.7359577 |
|  | MZB1 | 0.03355595 | -0.97666 | 21 | 0.79895123 |
|  | TNFRSF13B | 0.03554675 | -0.7473187 | 22 | 0.80788074 |
|  | NR4A1 | 0.03913421 | -0.7219743 | 23 | 0.85074369 |
|  | IGFBP3 | 0.0394714 | -0.8169124 | 24 | 0.82232089 |
|  | IGHG3 | 0.04344631 | -0.790011 | 25 | 0.86892625 |
|  | POU2AF1 | 0.04552957 | -0.6751142 | 26 | 0.87556871 |
|  | STRBP | 0.04823207 | -0.7163713 | 27 | 0.89318645 |

**Supplemental Table 4**

Regional differential expression – Lymphoid Follicles (pLN)

|  | gene | p_val | avg_log2FC | rank | BH.FDR |
| --- | --- | --- | --- | --- | --- |
| **Lymphoid Follicle - T1D vs ND** | IGKC | 4.96E-42 | -0.9184021 | 1 | 2.48E-39 |
|  | LTB | 4.69E-10 | **1.19521424** | 2 | 1.17E-07 |
|  | IGHA1 | 5.95E-09 | **-1.1006557** | 3 | 9.92E-07 |
|  | JCHAIN | 6.93E-06 | **-1.1616839** | 4 | 0.0008659 |
|  | CLU | 3.38E-05 | **1.12720485** | 5 | 0.00338249 |
|  | NIBAN3 | 3.99E-05 | **1.02506755** | 6 | 0.00332418 |
|  | FDCSP | 5.53E-05 | **1.10815829** | 7 | 0.00394865 |
|  | FCER2 | 5.78E-05 | **1.00748747** | 8 | 0.003614 |
|  | IGHG1 | 0.00016911 | -0.5747823 | 9 | 0.00939519 |
|  | CD37 | 0.00083429 | 0.54560787 | 10 | 0.04171458 |
|  | MS4A1 | 0.00172527 | 0.51070443 | 11 | 0.07842149 |
|  | CD22 | 0.00212582 | 0.7115422 | 12 | 0.08857592 |
|  | CD79A | 0.00372287 | 0.52904534 | 13 | 0.14318732 |
|  | CCL21 | 0.00450254 | 0.55852251 | 14 | 0.16080491 |
|  | TXNDC5 | 0.00498703 | -0.9232772 | 15 | 0.16623438 |
|  | CD72 | 0.00667488 | 0.96585187 | 16 | 0.20859013 |
|  | CCL19 | 0.01182727 | 0.89037932 | 17 | 0.34786084 |
|  | C3 | 0.01513201 | 0.89428554 | 18 | 0.42033371 |
|  | IGHD | 0.01903111 | 0.61944262 | 19 | 0.50081873 |
|  | CD79B | 0.02329492 | 0.57675527 | 20 | 0.58237302 |
|  | MYO7B | 0.02392616 | 1.0162796 | 21 | 0.5696704 |
|  | IGLC1 | 0.02452851 | -0.8200603 | 22 | 0.55746612 |
|  | TCL1A | 0.03032059 | 0.85377081 | 23 | 0.65914327 |
|  | SMIM14 | 0.03415679 | 0.70840607 | 24 | 0.71159974 |
|  | VPREB3 | 0.03793275 | 0.72524818 | 25 | 0.75865495 |

|  | gene | p_val | avg_log2FC | rank | BH.FDR |
| --- | --- | --- | --- | --- | --- |
| **Lymphoid Follicle - T1D vs AAb+** | IGHM | 2.69E-14 | 0.97741733 | 1 | 1.34E-11 |
|  | IGKC | 3.98E-14 | 0.6825293 | 2 | 9.94E-12 |
|  | LTB | 1.22E-12 | **1.46224594** | 3 | 2.04E-10 |
|  | CD79A | 6.79E-09 | **1.25775052** | 4 | 8.49E-07 |
|  | CD37 | 7.15E-09 | **1.06761842** | 5 | 7.15E-07 |
|  | CD52 | 8.77E-07 | **1.2254938** | 6 | 7.31E-05 |
|  | P2RX5 | 1.22E-06 | **1.29204526** | 7 | 8.70E-05 |
|  | CCL21 | 5.06E-06 | **0.99143182** | 8 | 0.00031638 |
|  | MS4A1 | 5.97E-06 | 0.78446512 | 9 | 0.00033194 |
|  | CD79B | 1.56E-05 | **1.31301846** | 10 | 0.00077873 |
|  | BANK1 | 2.51E-05 | **1.13860654** | 11 | 0.00114082 |
|  | TNFRSF13C | 5.53E-05 | **1.02166347** | 12 | 0.00230338 |
|  | CD22 | 0.00079496 | 0.79129459 | 13 | 0.03057535 |
|  | POU2AF1 | 0.00080655 | **1.07040936** | 14 | 0.02880551 |
|  | STRBP | 0.00109722 | **1.1269031** | 15 | 0.03657397 |
|  | CR1 | 0.00223318 | 1.05386026 | 16 | 0.0697869 |
|  | VPREB3 | 0.00269912 | 1.16945128 | 17 | 0.07938591 |
|  | NIBAN3 | 0.00334654 | 0.6739753 | 18 | 0.09295936 |
|  | FDCSP | 0.00360674 | 1.36159404 | 19 | 0.09491416 |
|  | SPIB | 0.00362279 | 1.05951701 | 20 | 0.0905698 |
|  | TNFRSF13B | 0.00503974 | 0.95715076 | 21 | 0.11999386 |
|  | SMIM14 | 0.00545499 | 0.99407022 | 22 | 0.12397694 |
|  | C3 | 0.00547922 | 1.06781343 | 23 | 0.1191134 |
|  | STK17B | 0.00598227 | 0.83342225 | 24 | 0.12463069 |
|  | FOSB | 0.00691896 | -1.2691026 | 25 | 0.13837916 |
|  | FCMR | 0.00737365 | 0.7536368 | 26 | 0.14180091 |
|  | PAX5 | 0.00784042 | 0.93362191 | 27 | 0.14519303 |
|  | FCRL1 | 0.00864951 | 0.90321148 | 28 | 0.15445561 |
|  | CCL19 | 0.00889728 | 0.93542199 | 29 | 0.15340138 |
|  | BLK | 0.00978362 | 0.65535277 | 30 | 0.16306029 |
|  | FCRL2 | 0.00990895 | 0.88474207 | 31 | 0.15982169 |
|  | CXCR5 | 0.01031539 | 1.04550562 | 32 | 0.16117789 |
|  | TLR10 | 0.011076 | 0.92885396 | 33 | 0.16781824 |
|  | CLEC4G | 0.01138048 | -1.1082241 | 34 | 0.16736004 |
|  | IGHA1 | 0.01243627 | 0.62648788 | 35 | 0.17766097 |
|  | TMC8 | 0.01563927 | 0.66079666 | 36 | 0.21721213 |
|  | STAB2 | 0.01698737 | -1.1544503 | 37 | 0.22955904 |
|  | BCL11A | 0.0172472 | 0.77593462 | 38 | 0.2269368 |
|  | FCRL3 | 0.01742589 | 0.95812593 | 39 | 0.22340887 |
|  | FCER2 | 0.02005419 | 0.89602701 | 40 | 0.2506774 |
|  | FCRLA | 0.02209493 | 1.02023571 | 41 | 0.26945033 |
|  | TAF11L6 | 0.02315792 | -1.1465418 | 42 | 0.27568955 |
|  | WDFY4 | 0.02373114 | 0.82677905 | 43 | 0.27594351 |
|  | ARHGAP15 | 0.02737151 | 0.63003309 | 44 | 0.31103984 |
|  | MMP19 | 0.02857103 | -1.0185111 | 45 | 0.31745587 |
|  | CD72 | 0.02869322 | 0.7382461 | 46 | 0.31188279 |
|  | FOXD4L5 | 0.02874561 | -1.0425773 | 47 | 0.30580437 |
|  | CD19 | 0.02919172 | 0.77850807 | 48 | 0.30408044 |
|  | IGHD | 0.03303574 | 0.55535001 | 49 | 0.33709937 |
|  | CLU | 0.03580789 | 0.49228324 | 50 | 0.35807889 |
|  | MYO7B | 0.04283672 | 0.8793646 | 51 | 0.41996786 |
|  | IGHG1 | 0.04285747 | 0.36522876 | 52 | 0.41209108 |
|  | ADAM28 | 0.04346795 | 0.65738677 | 53 | 0.41007502 |
|  | CD1C | 0.04457877 | 0.92960946 | 54 | 0.41276635 |
|  | CR2 | 0.04813643 | 0.81580468 | 55 | 0.43760393 |
