## Supplemental Table 8 for "Spatial transcriptomics from pancreas and local draining lymph node tissue reveals a lymphotoxin-β signature in human type 1 diabetes"

**Supplemental Table 5**

Regional differential expression – T-cell Zone (pLN)

|  | gene | p_val | avg_log2FC | rank | BH.FDR |
| --- | --- | --- | --- | --- | --- |
| **T-cell Zone - AAb+ vs ND** | IGKC | 2.77E-43 | **-1.2555638** | 1 | 1.38E-40 |
|  | IGHM | 8.29E-12 | **-1.3354532** | 2 | 2.07E-09 |
|  | IGHA1 | 1.86E-09 | **-1.2979191** | 3 | 3.10E-07 |
|  | CCL19 | 0.00639744 | 0.76662347 | 4 | 0.79967952 |
|  | TNIK | 0.00660569 | 0.95011041 | 5 | 0.66056874 |
|  | TRAC | 0.00689551 | 0.57458653 | 6 | 0.57462616 |
|  | ZAP70 | 0.00886145 | 0.95219725 | 7 | 0.63296043 |
|  | CHD3 | 0.01137267 | 0.90547161 | 8 | 0.71079166 |
|  | TRBC1 | 0.01353958 | 0.66687178 | 9 | 0.75219912 |
|  | CCN1 | 0.01466463 | -0.8930835 | 10 | 0.73323162 |
|  | JAK3 | 0.01750282 | 0.84311734 | 11 | 0.79558255 |
|  | TBC1D10C | 0.01893461 | 0.72096443 | 12 | 0.78894217 |
|  | LTB | 0.01914547 | 0.59965815 | 13 | 0.73636428 |
|  | FAM118A | 0.02023371 | 1.03809019 | 14 | 0.72263264 |
|  | FOSB | 0.02202468 | 1.05141679 | 15 | 0.73415592 |
|  | GIMAP7 | 0.02222906 | 0.69513685 | 16 | 0.69465807 |
|  | NLRC5 | 0.02281245 | 0.79104783 | 17 | 0.67095447 |
|  | MYL9 | 0.02851717 | -0.9046784 | 18 | 0.79214354 |
|  | TRBC2 | 0.02857203 | 0.49461069 | 19 | 0.75189543 |
|  | DGKA | 0.0317319 | 0.64281646 | 20 | 0.79329744 |
|  | TXNDC5 | 0.03351816 | -0.8286621 | 21 | 0.79805132 |
|  | IGHG1 | 0.03590645 | -0.3581218 | 22 | 0.81605567 |
|  | OXNAD1 | 0.03627909 | 0.67288221 | 23 | 0.78867596 |
|  | EVL | 0.04253271 | 0.50521539 | 24 | 0.88609821 |

|  | gene | p_val | avg_log2FC | rank | BH.FDR |
| --- | --- | --- | --- | --- | --- |
| **T-cell Zone - T1D vs ND** | IGKC | 1.23E-22 | -0.7503586 | 1 | 6.17E-20 |
|  | IGHA1 | 2.14E-08 | **-1.0793471** | 2 | 5.36E-06 |
|  | CCL19 | 2.96E-05 | **1.11004225** | 3 | 0.00492657 |
|  | IGHM | 0.0004463 | -0.5441709 | 4 | 0.05578723 |
|  | IGLC1 | 0.00585522 | -1.0423039 | 5 | 0.58552231 |
|  | JCHAIN | 0.0078349 | -0.6939855 | 6 | 0.65290824 |
|  | C3 | 0.01195238 | 0.74122764 | 7 | 0.85374137 |
|  | CCL21 | 0.0128535 | 0.35515187 | 8 | 0.80334361 |
|  | SATB1 | 0.03221505 | -0.8205765 | 9 | 1.78972502 |
|  | AQP1 | 0.0334492 | 0.9713346 | 10 | 1.67246013 |
|  | FCMR | 0.04279399 | -0.6618083 | 11 | 1.94518141 |
|  | IGHG1 | 0.04538095 | -0.3231766 | 12 | 1.890873 |
|  | MMP12 | 0.04785785 | 1.46378593 | 13 | 1.84068668 |

|  | gene | p_val | avg_log2FC | rank | BH.FDR |
| --- | --- | --- | --- | --- | --- |
| **T-cell Zone - T1D vs AAb+** | IGKC | 1.45E-07 | 0.50520517 | 1 | 7.25E-05 |
|  | IGHM | 7.53E-05 | 0.79128226 | 2 | 0.01882129 |
|  | CCL21 | 0.00017771 | 0.55892035 | 3 | 0.02961742 |
|  | CLEC2D | 0.00125315 | -1.054121 | 4 | 0.15664378 |
|  | SATB1 | 0.00287368 | -1.1259035 | 5 | 0.28736838 |
|  | FOSB | 0.00331013 | -1.3863774 | 6 | 0.27584389 |
|  | EVL | 0.00410994 | -0.7039052 | 7 | 0.29356702 |
|  | DGKA | 0.0094434 | -0.7504285 | 8 | 0.59021277 |
|  | TRBC2 | 0.00977567 | -0.5609361 | 9 | 0.54309264 |
|  | TNIK | 0.01170511 | -0.8058644 | 10 | 0.58525556 |
|  | FYB1 | 0.01399239 | -0.7558621 | 11 | 0.63601763 |
|  | NELL2 | 0.01428275 | -1.061959 | 12 | 0.59511438 |
|  | TRAC | 0.01622685 | -0.4761481 | 13 | 0.62410946 |
|  | TMC8 | 0.0199656 | -0.582903 | 14 | 0.71305712 |
|  | TRBC1 | 0.02017009 | -0.5848705 | 15 | 0.6723362 |
|  | LEF1 | 0.02024986 | -0.8468521 | 16 | 0.63280814 |
|  | S100A9 | 0.02789269 | 1.07515644 | 17 | 0.82037325 |
|  | FOXD4L5 | 0.02829736 | -1.1031134 | 18 | 0.78603786 |
|  | ARHGAP15 | 0.02907046 | -0.6765752 | 19 | 0.76501209 |
|  | MMP12 | 0.02964293 | 1.19820981 | 20 | 0.74107326 |
|  | GIMAP5 | 0.0306555 | -0.818942 | 21 | 0.72989278 |
|  | FOS | 0.03080969 | -0.7872021 | 22 | 0.70022019 |
|  | CHD3 | 0.03090074 | -0.6981739 | 23 | 0.67175513 |
|  | STK17B | 0.0324315 | -0.6801146 | 24 | 0.67565619 |
|  | MMP19 | 0.03280442 | -0.9820746 | 25 | 0.65608841 |
|  | AQP1 | 0.03545067 | 0.95683213 | 26 | 0.68174369 |
|  | TCF7 | 0.03554229 | -0.5802262 | 27 | 0.6581905 |
|  | TAF11L6 | 0.04137761 | -1.0134762 | 28 | 0.73888584 |
|  | STK17A | 0.04258047 | -0.8131519 | 29 | 0.73414598 |
|  | CD3E | 0.0449233 | -0.639932 | 30 | 0.74872167 |
|  | JAK3 | 0.0490875 | -0.6325067 | 31 | 0.79173389 |
