## Supplemental Table 9 for "Spatial transcriptomics from pancreas and local draining lymph node tissue reveals a lymphotoxin-β signature in human type 1 diabetes"

**Supplemental Table 6**

Regional differential expression – Stroma (pLN)

|  | gene | p_val | avg_log2FC | rank | BH.FDR |
| --- | --- | --- | --- | --- | --- |
| **Stroma - AAb+ vs ND** | IGKC | 7.06E-73 | **-1.4363382** | 1 | 3.53E-70 |
|  | IGHG1 | 6.92E-11 | -0.9525089 | 2 | 1.73E-08 |
|  | IGHA1 | 8.13E-11 | **-1.2616625** | 3 | 1.35E-08 |
|  | IGHM | 1.87E-10 | **-1.030625** | 4 | 2.34E-08 |
|  | TXNDC5 | 0.00164813 | -1.2294228 | 5 | 0.16481305 |
|  | CCN1 | 0.00326165 | -1.111371 | 6 | 0.27180374 |
|  | PLTP | 0.00357402 | -1.0944894 | 7 | 0.25528697 |
|  | MYL9 | 0.00588949 | -0.7227732 | 8 | 0.36809329 |
|  | IGLC1 | 0.0067283 | -0.957743 | 9 | 0.37379457 |
|  | S100A9 | 0.01089763 | -1.3210835 | 10 | 0.54488164 |
|  | JCHAIN | 0.01636832 | -0.5984099 | 11 | 0.7440146 |
|  | RNASE1 | 0.01808997 | -0.8656722 | 12 | 0.75374881 |
|  | EGR1 | 0.02677612 | -0.6565103 | 13 | 1.02985079 |
|  | LYVE1 | 0.03228843 | -0.861776 | 14 | 1.1531583 |
|  | MZB1 | 0.03685289 | -1.0282333 | 15 | 1.22842957 |

|  | gene | p_val | avg_log2FC | rank | BH.FDR |
| --- | --- | --- | --- | --- | --- |
| **Stroma - T1D vs ND** | IGKC | 3.66E-80 | **-1.4015655** | 1 | 1.83E-77 |
|  | IGHM | 1.20E-18 | **-1.4961667** | 2 | 3.00E-16 |
|  | IGHA1 | 2.79E-14 | **-1.4499387** | 3 | 4.65E-12 |
|  | IGHG1 | 2.57E-12 | -0.9670311 | 4 | 3.21E-10 |
|  | IGLC1 | 0.00027478 | **-1.3113685** | 5 | 0.02747803 |
|  | JCHAIN | 0.00029634 | -0.9084944 | 6 | 0.02469504 |
|  | C3 | 0.00083837 | 1.0303635 | 7 | 0.05988381 |
|  | DES | 0.0009991 | 0.88805088 | 8 | 0.06244347 |
|  | CCN1 | 0.00180622 | -1.1035761 | 9 | 0.10034555 |
|  | FABP4 | 0.00237519 | 1.19075173 | 10 | 0.11875949 |
|  | TXNDC5 | 0.00271974 | -1.0519888 | 11 | 0.12362432 |
|  | FASN | 0.00482632 | 1.451 | 12 | 0.20109651 |
|  | FOSB | 0.00839538 | -1.1239631 | 13 | 0.3228993 |
|  | RNASE1 | 0.01360437 | 0.68803485 | 14 | 0.48587038 |
|  | PLA2G2A | 0.02192155 | 1.07555022 | 15 | 0.73071848 |
|  | CD5L | 0.0242326 | -1.1200848 | 16 | 0.75726877 |
|  | MMP19 | 0.02455333 | -1.0413207 | 17 | 0.72215662 |
|  | CCDC80 | 0.02628886 | 1.09277316 | 18 | 0.73024601 |
|  | MZB1 | 0.03926195 | -0.9236982 | 19 | 1.03320908 |
|  | SCD | 0.03967524 | 1.04072835 | 20 | 0.99188095 |

|  | gene | p_val | avg_log2FC | rank | BH.FDR |
| --- | --- | --- | --- | --- | --- |
| **Stroma - T1D vs AAb+** | RNASE1 | 8.17E-06 | **1.55370709** | 1 | 0.00408263 |
|  | PLTP | 1.61E-05 | **1.56582273** | 2 | 0.00401419 |
|  | FOSB | 6.13E-05 | **-1.722657** | 3 | 0.01021911 |
|  | MYL9 | 0.00016621 | 0.94015346 | 4 | 0.0207756 |
|  | C3 | 0.0002259 | **1.18314763** | 5 | 0.0225904 |
|  | S100A9 | 0.00035771 | **1.94850577** | 6 | 0.02980882 |
|  | DES | 0.00041907 | 0.97131227 | 7 | 0.02993338 |
|  | F13A1 | 0.00378078 | 1.3365791 | 8 | 0.23629846 |
|  | FABP4 | 0.0039264 | 1.10412433 | 9 | 0.21813323 |
|  | PLA2G2A | 0.00815735 | 1.39039749 | 10 | 0.40786772 |
|  | MMP19 | 0.0089675 | -1.2283652 | 11 | 0.40761349 |
|  | FASN | 0.01203242 | 1.16992864 | 12 | 0.50135064 |
|  | TAF11L6 | 0.01318003 | -1.524119 | 13 | 0.50692421 |
|  | IGHM | 0.01528828 | -0.4655418 | 14 | 0.54601007 |
|  | S100A8 | 0.01653343 | 1.32609296 | 15 | 0.55111438 |
|  | USP17L15 | 0.01669517 | -1.2882201 | 16 | 0.52172395 |
|  | FOXD4L5 | 0.01926733 | -1.3789921 | 17 | 0.56668616 |
|  | USP17L3 | 0.02180815 | -1.2021325 | 18 | 0.60578192 |
|  | SCD | 0.03196735 | 1.12135816 | 19 | 0.84124613 |
|  | CLEC4M | 0.03549935 | -0.9601693 | 20 | 0.88748365 |
|  | CCDC80 | 0.03673249 | 0.99627672 | 21 | 0.87458316 |
|  | TAF11L8 | 0.04041286 | -1.1721767 | 22 | 0.91847397 |
