## Supplemental Table 10 for "Spatial transcriptomics from pancreas and local draining lymph node tissue reveals a lymphotoxin-β signature in human type 1 diabetes"

**Supplemental Table 7**

Regional differential expression – Medulla (pLN)

|  | gene | p_val | avg_log2FC | rank | BH.FDR |
| --- | --- | --- | --- | --- | --- |
| **Medulla - AAb+ vs ND** | IGKC | 1.16E-93 | **-1.5747661** | 1 | 5.81E-91 |
|  | IGHA1 | 5.95E-14 | **-1.8701375** | 2 | 1.49E-11 |
|  | IGHG1 | 5.87E-13 | **-1.14944** | 3 | 9.78E-11 |
|  | IGHM | 2.00E-10 | -0.8700865 | 4 | 2.49E-08 |
|  | STAB2 | 7.98E-05 | 0.92608685 | 5 | 0.00797714 |
|  | JCHAIN | 0.00011752 | **-1.022461** | 6 | 0.00979339 |
|  | FOSB | 0.00019016 | **1.706681** | 7 | 0.01358296 |
|  | TXNDC5 | 0.00033047 | **-1.3761096** | 8 | 0.02065457 |
|  | IGLC1 | 0.0003623 | **-1.4078176** | 9 | 0.02012764 |
|  | PLTP | 0.00065366 | **-1.1048563** | 10 | 0.0326831 |
|  | CLEC4M | 0.00111898 | 0.79502014 | 11 | 0.05086278 |
|  | CLEC4G | 0.00228621 | 0.58388605 | 12 | 0.09525887 |
|  | S100A9 | 0.00413437 | -1.2884319 | 13 | 0.15901408 |
|  | EGR1 | 0.00433641 | -0.8948103 | 14 | 0.15487166 |
|  | LMNA | 0.00481993 | 0.9847971 | 15 | 0.16066447 |
|  | CCN1 | 0.0062312 | -1.2620094 | 16 | 0.19472492 |
|  | IGHG3 | 0.00968591 | -0.9524412 | 17 | 0.28487965 |
|  | CD52 | 0.01136658 | -1.1111405 | 18 | 0.31573835 |
|  | CCL21 | 0.01212519 | -0.8143396 | 19 | 0.31908406 |
|  | MARCO | 0.0145727 | 0.66911322 | 20 | 0.36431738 |
|  | MZB1 | 0.01794872 | -1.1415157 | 21 | 0.42735051 |
|  | MMP19 | 0.01859341 | 0.91851308 | 22 | 0.42257754 |
|  | LGMN | 0.02082987 | 0.67418074 | 23 | 0.45282329 |
|  | FOS | 0.03578655 | 0.63023183 | 24 | 0.74555312 |
|  | CXCL2 | 0.03885017 | -0.7372649 | 25 | 0.77700331 |

|  | gene | p_val | avg_log2FC | rank | BH.FDR |
| --- | --- | --- | --- | --- | --- |
| **Medulla - T1D vs ND** | IGKC | 8.89E-54 | -0.961279 | 1 | 4.44E-51 |
|  | IGHM | 5.05E-10 | -0.7899031 | 2 | 1.26E-07 |
|  | IGHG1 | 1.10E-06 | -0.6596627 | 3 | 0.00018377 |
|  | IGHA1 | 3.72E-06 | -0.8388632 | 4 | 0.00046482 |
|  | MARCO | 8.59E-05 | **1.01262174** | 5 | 0.00859465 |
|  | CLEC4G | 0.00038137 | 0.65232324 | 6 | 0.03178085 |
|  | IGLC1 | 0.00069713 | **-1.1815742** | 7 | 0.04979529 |
|  | CXCL2 | 0.00075437 | **-1.2755273** | 8 | 0.04714791 |
|  | JCHAIN | 0.00081617 | -0.798288 | 9 | 0.0453426 |
|  | LYZ | 0.00175384 | 0.77986073 | 10 | 0.0876918 |
|  | RNASE1 | 0.00327886 | 0.82158169 | 11 | 0.14903895 |
|  | IGHG3 | 0.01325072 | -0.8379483 | 12 | 0.5521135 |
|  | CLU | 0.01443453 | 0.7986649 | 13 | 0.55517415 |
|  | CCL19 | 0.02416584 | 0.88238904 | 14 | 0.86306561 |
|  | TXNDC5 | 0.02715243 | -0.6796452 | 15 | 0.90508098 |
|  | FABP4 | 0.02899806 | 0.8883583 | 16 | 0.90618932 |
|  | LGMN | 0.04088021 | 0.58242781 | 17 | 1.2023592 |
|  | SPP1 | 0.04341409 | 2.21962217 | 18 | 1.20594685 |
|  | LTB | 0.04654457 | 0.64129548 | 19 | 1.22485717 |

|  | gene | p_val | avg_log2FC | rank | BH.FDR |
| --- | --- | --- | --- | --- | --- |
| **Medulla - T1D vs AAb+** | IGKC | 9.20E-14 | 0.61348706 | 1 | 4.60E-11 |
|  | FOSB | 2.02E-05 | **-2.2379644** | 2 | 0.00505639 |
|  | S100A9 | 4.93E-05 | **1.81881737** | 3 | 0.0082149 |
|  | IGHA1 | 5.91E-05 | **1.03127439** | 4 | 0.00738524 |
|  | RNASE1 | 6.66E-05 | **1.23513419** | 5 | 0.00665722 |
|  | PLTP | 0.00017835 | **1.18054211** | 6 | 0.01486247 |
|  | CCL21 | 0.00044189 | **1.08954723** | 7 | 0.03156348 |
|  | FOS | 0.000823 | -1.0296305 | 8 | 0.05143761 |
|  | CLU | 0.00127121 | 1.14611951 | 9 | 0.07062254 |
|  | MMP19 | 0.00233331 | -1.1858033 | 10 | 0.11666547 |
|  | LTB | 0.00253788 | 1.07861199 | 11 | 0.11535823 |
|  | FABP4 | 0.00264844 | 1.43174764 | 12 | 0.11035172 |
|  | SPP1 | 0.0027383 | 1.9325628 | 13 | 0.10531917 |
|  | IGHG1 | 0.00322468 | 0.48977727 | 14 | 0.11516728 |
|  | LMNA | 0.00338908 | -0.9570293 | 15 | 0.11296927 |
|  | CCL19 | 0.00732062 | 1.11373856 | 16 | 0.22876921 |
|  | CD52 | 0.01152563 | 1.08487528 | 17 | 0.33898902 |
|  | FSTL3 | 0.01379808 | -1.1207574 | 18 | 0.38328008 |
|  | CD5L | 0.01525881 | 0.87540892 | 19 | 0.40154775 |
|  | F13A1 | 0.01537446 | 1.13582361 | 20 | 0.38436151 |
|  | STAB2 | 0.01718561 | -0.4839197 | 21 | 0.40918127 |
|  | FOXD4L5 | 0.02088734 | -1.2450628 | 22 | 0.47471233 |
|  | TAF11L6 | 0.02174271 | -1.2267142 | 23 | 0.47266751 |
|  | SERPINE1 | 0.02230611 | -1.1070748 | 24 | 0.46471064 |
|  | CD37 | 0.0277011 | 0.77195164 | 25 | 0.55402196 |
|  | NTS | 0.02886808 | 1.09827011 | 26 | 0.55515542 |
|  | P2RX5 | 0.03094314 | 0.96216351 | 27 | 0.57302113 |
|  | S100A8 | 0.03293232 | 1.03573736 | 28 | 0.58807718 |
|  | HIST1H1B | 0.03758877 | 0.97026845 | 29 | 0.64808227 |
|  | C3 | 0.04053702 | 0.90025318 | 30 | 0.67561697 |
|  | CLEC4M | 0.04079161 | -0.4412128 | 31 | 0.65792923 |
|  | USP17L3 | 0.04928958 | -0.9716744 | 32 | 0.77014969 |
|  | ACKR1 | 0.04942316 | 0.89625258 | 33 | 0.74883576 |
